# Digit ratio (2D:4D) and Congenital Adrenal Hyperplasia (CAH): Systematic Literature Review and Meta-Analysis

**DOI:** 10.1101/2020.05.27.119115

**Authors:** Gareth Richards, Wendy V. Browne, Ezra Aydin, Mihaela Constantinescu, Gideon Nave, Mimi S. Kim, Steven J. Watson

## Abstract

The ratio of length between the second and fourth fingers (2D:4D) is commonly used as an indicator of prenatal sex hormone exposure. Several approaches have been used to try to validate the measure, including examining 2D:4D in people with congenital adrenal hyperplasia (CAH), a suite of conditions characterised by elevated adrenal androgen production secondary to defective steroidogenesis. We present here a systematic review that examines the relationship between these two variables. Twelve articles relating to nine CAH cohorts were identified, and 2D:4D comparisons have been made between cases and controls in eight of these cohorts. Altogether, at least one 2D:4D variable has been compared between n=251 females with CAH and n=358 unaffected females, and between n=108 males with CAH and n=204 unaffected males. A previous meta-analysis (Hönekopp & Watson, 2010) reported lower right hand (R2D:4D) and left hand (L2D:4D) digit ratios in patients with CAH relative to sex-matched controls. Our meta-analysis showed the same direction of results; however, the effects were only statistically significant for R2D:4D in males and L2D:4D in females (R2D:4D: females, *p* = 0.072, *g* = 0.591; males, *p* = 0.019, *g* = 0.513; L2D:4D: females, *p* = 0.020, *g* = 0.245; males, *p* = 0.334, *g* = 0.218), and the average effect size had reduced by 46.70%. We also found no evidence to suggest the right-left difference in 2D:4D (D_[R-L]_) is associated with prenatal sex hormone exposure.

## Introduction

Digit ratio (2D:4D) is typically lower in males than females, with a slightly larger sex difference present for the right hand (Hönekopp & Watson, 2010). The measure is frequently used as an indicator of the effects of prenatal sex hormones, and has been suggested to index the level of exposure to foetal testosterone (Brown, Hines, Fane, & Breedlove, 2002; Manning, Scutt, Wilson, & Lewis-Jones, 1998) or the ratio of foetal testosterone to foetal oestradiol (Lutchmaya, Baron-Cohen, Raggatt, Knickmeyer, & Manning, 2004; Manning, 2011; Zheng & Cohn, 2011). Although 2D:4D is commonly used to examine the effects of the prenatal hormonal environment on later phenotype, relatively few studies have attempted to validate the measure in human populations.

Some research has directly manipulated foetal hormones in animal models (Abbott, Colman, Tiefenthaler, Dumesic, & Abbott, 2012; Auger et al., 2013; Huber, Lenz, Kornhuber, & Müller, 2017; Romano, Rubolini, Martinelli, Alquati, & Saino, 2005; Saino, Rubolini, Romano, & Boncoraglio, 2007; Talarovičová, Kršková, & Blažeková, 2009; Zheng & Cohn, 2011), though the effects reported have not always been consistent. For instance, although Zheng and Cohn (2011) and Huber et al. (2017) both examined the effects of prenatal hormone exposure in CD-1 mice, the studies reported effects in opposing directions. Early manipulation of hormones is unethical in human studies, meaning that researchers have had to rely on other methods, such as correlating 2D:4D with hormone concentrations in amniotic fluid (Lutchmaya et al., 2004; Richards, Gomes, & Ventura, 2018; Ventura, Gomes, Pita, Neto, & Taylor, 2013), umbilical cord blood (Çetin, Can, & Özcan, 2016; Hickey et al., 2010; Hollier et al., 2015; Mitsui et al., 2016, 2015; Whitehouse et al., 2015), or the maternal circulation (Barona, Kothari, Skuse, & Micali, 2015; Hickey et al., 2010; Richards et al., 2018; Ventura et al., 2013). The results of studies in humans broadly point toward a negative correlation between foetal testosterone exposure and 2D:4D, although the statistically significant effects are accompanied by many null findings (Richards, 2017), and publication bias may be an issue.

Another approach for determining the efficacy of 2D:4D has been to examine whether it is associated with medical conditions characterised by atypical androgen activity. Two studies (Berenbaum, Bryk, Nowak, Quigley, & Moffat, 2009; van Hemmen, Cohen-Kettenis, Steensma, Veltman, & Bakker, 2017) have reported evidence of feminised 2D:4D ratios in phenotypically female (46XY) individuals with complete androgen insensitivity syndrome (CAIS), although it should be noted that the variance for 2D:4D in this population appears to be comparable to that of controls despite the complete lack of androgen sensitivity (Berenbaum et al., 2009; see also commentary by Wallen, 2009). Manning, Kilduff, and Trivers (2013) showed that digit ratios were higher (i.e. more female-typical) in males with Klinefelter syndrome (47XXY) than in their unaffected relatives. However, this effect is difficult to interpret considering that prenatal testosterone levels in males with Klinefelter syndrome do not appear to differ from those of typically developing males (Ratcliffe et al., 1994).

A promising area of research has examined individuals with congenital adrenal hyperplasia (CAH). CAH is a family of autosomal recessive conditions characterised by impairment of one of five enzymes required to synthesise cortisol from cholesterol. This causes an accumulation of adrenocorticotrophic hormone (ACTH) secondary to negative feedback, which results in overstimulation of the adrenal cortex and increased adrenal androgen production (New, 2006). Most cases (90–95%) of CAH are caused by 21–hydroxylase (21–OH) deficiency, with three main phenotypes being distinguishable (for a comparison of symptom profiles, see New, 2006). The most severe form, classical salt-wasting (SW) CAH, involves impairment of aldosterone synthesis, a symptom that is absent overall in classical simple-virilizing (SV) CAH; both SW and SV are characterised by genital ambiguity in female (46XX) patients. Pharmacological treatment for classical CAH due to 21-OH deficiency typically begins soon after birth, and the condition has been found to occur in approximately 1 in 14,000 live births (Pang et al., 1988). Non-classical CAH due to 21-OH deficiency does not present with aldosterone impairment nor typically with genital ambiguity, and can go undetected (Levine et al., 1980) particularly in males. The non-classical or late-onset form is diagnosed when symptoms present later in life (Kisch, Laurian, & Hoerer, 1987; New, Dupont, Pollack, & Levine, 1981), and is more common than classical CAH, with reported prevalence ranging from 1 in 27 to 1 in 300, depending on the ethnic group studied (Hannah-Shmouni et al., 2017; New, 2006).

CAH provides an opportunity for researchers to examine the organisational effects of elevated androgen exposure during gestation. There is some evidence for behavioural masculinisation and defeminisation in CAH, with such issues being particularly pertinent in 46XX female-assigned cases because prenatal androgen concentrations may not only affect external somatic sex structures, but also bipotential areas in the brain, leading to modification of behavioural/psychological outcomes (see Cohen-Bendahan, van de Beek, & Berenbaum, 2005; Hines, 2004; Hines, Constantinescu, & Spencer, 2015; Jordan-Young, 2012). The early androgenic effects of CAH in males however are less clear, as feedback mechanisms may lead to normalisation of androgen levels via reduced production by the testes (Pang, Levine, Chow, Faiman, & New, 1979; for a discussion, see Mathews et al., 2004). Evidence for this is provided by the observation that amniotic testosterone levels for 46XY CAH foetuses tend not to be clearly distinguishable from those of typically developing 46XY foetuses (Pang et al., 1980; Wudy, Dörr, Solleder, Djalali, & Homoki, 1999), though such observations have relied on very small samples. However, it does appear possible that following an initial elevation, testosterone concentrations may be relatively normal in males with CAH.

Hines et al. (2003) reported that females with CAH outperformed their unaffected female relatives on two tasks assessing targeting performance. The tasks employed included measures of visuomotor spatial ability that have been found to demonstrate a large male advantage in the typically-developing population (Watson, 2001). In a study by Collaer, Brook, Conway, Hindmarsh, and Hines (2009), females with CAH also outperformed unaffected female relatives on motor and visuomotor tasks (grip strength and targeting), which have shown a male advantage in previous research (e.g. Miller, MacDougall, Tarnopolsky, & Sale, 1993), even after controlling for weight and height. The enhanced targeting performance in females with CAH was still present after adjusting for grip strength, leading the researchers to point towards an organisational influence of prenatal androgens on the neural regions dedicated to targeting (Collaer et al., 2009).

Behavioural masculinisation in females with CAH may only occur in traits which show a particularly large male advantage. Alternatively, as studies of CAH populations typically utilise small samples due to the rarity of the condition, they may lack the statistical power required to reliably detect effects of small or medium size. A way to overcome this limitation is to pool the findings of individual studies into a meta-analysis, which provides an indication of the presence (or absence) of an effect as well as its size. Using this technique, Puts et al. (2008) reported that females with CAH display an advantage on spatial tasks, whereas males with CAH display a disadvantage. However, although a more recent meta-analysis (Collaer & Hines, 2020) including a larger number of samples replicated the finding of reduced overall spatial ability in males with CAH relative to males controls, it did not find any difference between female CAH cases and controls.

As CAH (at least in females) is associated with elevated prenatal androgen levels, and 2D:4D is hypothesised to indicate individual differences in foetal testosterone exposure, it follows that they should be related. A meta-analysis of early studies of CAH case-control studies (Hönekopp & Watson, 2010) showed that digit ratio for the right hand (R2D:4D) (*d* = 0.91, *p* < 0.001) and left hand (L2D:4D) (*d* = 0.75, *p* = 0.007) were significantly lower (i.e. more male-typical) in females with CAH relative to female controls; R2D:4D (*d* = 0.94, *p* = 0.061) and L2D:4D (*d* = 0.63, *p* = 0.013) were also lower in males with CAH relative to male controls, albeit the effect for R2D:4D was not statistically significant at the *p* < 0.05 level.

Although behavioural effects associated with CAH may be explainable by environmental influences (Hines et al., 2015; Jordan-Young, 2012) such as the presence and extent of genital virilisation, alterations in the way that parents, teachers and others interact with children with CAH, it seems unlikely that these could affect a person’s digit ratio. However, it should be acknowledged that although CAH research may indicate that elevated prenatal testosterone exposure causes physical differences, such as a masculinised 2D:4D ratio, these findings cannot necessarily be extrapolated to indicate a similar influence on the developing brain.

The current study aims to build on the earlier meta-analysis by Hönekopp and Watson (2010) by: (1) updating their analysis to include new studies, and (2) incorporating a full systematic review of the relevant literature. Hönekopp and Watson (2010) incorporated their analysis of the relationship between 2D:4D and CAH into an article with a much broader remit. Therefore, this literature has yet to be comprehensively reviewed. We also extend their analysis in other ways. As it has been suggested that the right-left difference in digit ratio (D[R-L]) can provide a further marker of prenatal sex hormone activity, with low R2D:4D relative to L2D:4D hypothesised to indicate high androgen exposure (Manning, 2002; Manning, Kilduff, Cook, Crewther, & Fink, 2014), this variable will also be considered in the current study. Furthermore, some studies report on the average 2D:4D across the right and left hands (M2D:4D). Because studies comparing digit ratios between patients with CAH and controls have not so far investigated D_[R-L]_ or M2D:4D, we contacted the authors of relevant papers to request the necessary data. We hypothesised that R2D:4D, L2D:4D, and D[R-L] would each be significantly lower in males and females with CAH relative to male and female controls, respectively. Although not initially considered in our pre-registration (see next section), we also hypothesised that M2D:4D would be significantly lower in males and females with CAH relative to male and female controls, respectively.

## Material and Methods

We pre-registered our review and analysis plan on the Open Science Framework (osf.io/n2hse) prior to beginning the research. Studies were considered eligible for inclusion where they made at least one comparison of 2D:4D between individuals with a diagnosis of any form of CAH with a control group. We made no limitations on year or language of publication. Studies were excluded where they did not report the statistics necessary to make a comparison between CAH and sex-matched controls, or if they did not report primary data.

We searched (keyword, title, and abstract; no publication date restrictions were imposed) Ovid MEDLINE, Embase, PsychINFO, Web of Science, and Scopus using the following search terms: (Digit ratio OR Digit length ratio OR Digital ratio OR Finger ratio OR Finger length ratio OR 2D:4D OR 2D4D OR Second to fourth OR Second-to-fourth OR Second-fourth OR 2nd to 4th OR 2nd-to-fourth OR 2nd-4th OR Ring to index OR Ring-to-index OR Index to ring OR Index-to-ring) AND (Congenital adrenal hyperplasia OR CAH). We also examined the reference lists of relevant papers, a bibliographic article of 2D:4D studies published between 1998 and 2008 (Voracek & Loibl, 2009), and an online database of digit ratio research (Fink & Manning, 2018), which (as of 09/12/2018) included 817 references. Additionally, we contacted 70+ researchers within the digit ratio and CAH fields to try to identify any published or unpublished data that we had not already included within our review.

We identified 3,705 articles through literature searches, four from reference lists of relevant papers, and three by contacting authors in the field. Once duplicates had been excluded, this resulted in 3,408 articles. The title for each was read, and the article was excluded from further consideration if it did not appear to relate to either 2D:4D or CAH. The abstracts of 615 potentially relevant articles were then accessed (please note that in cases where the article did not include an abstract, the Introduction, Introduction and Method, or whole article was read). Relevant materials that were not available in English were translated. See **Figure 1** for the PRISMA flow diagram (Moher et al., 2009).

**Figure 1.**
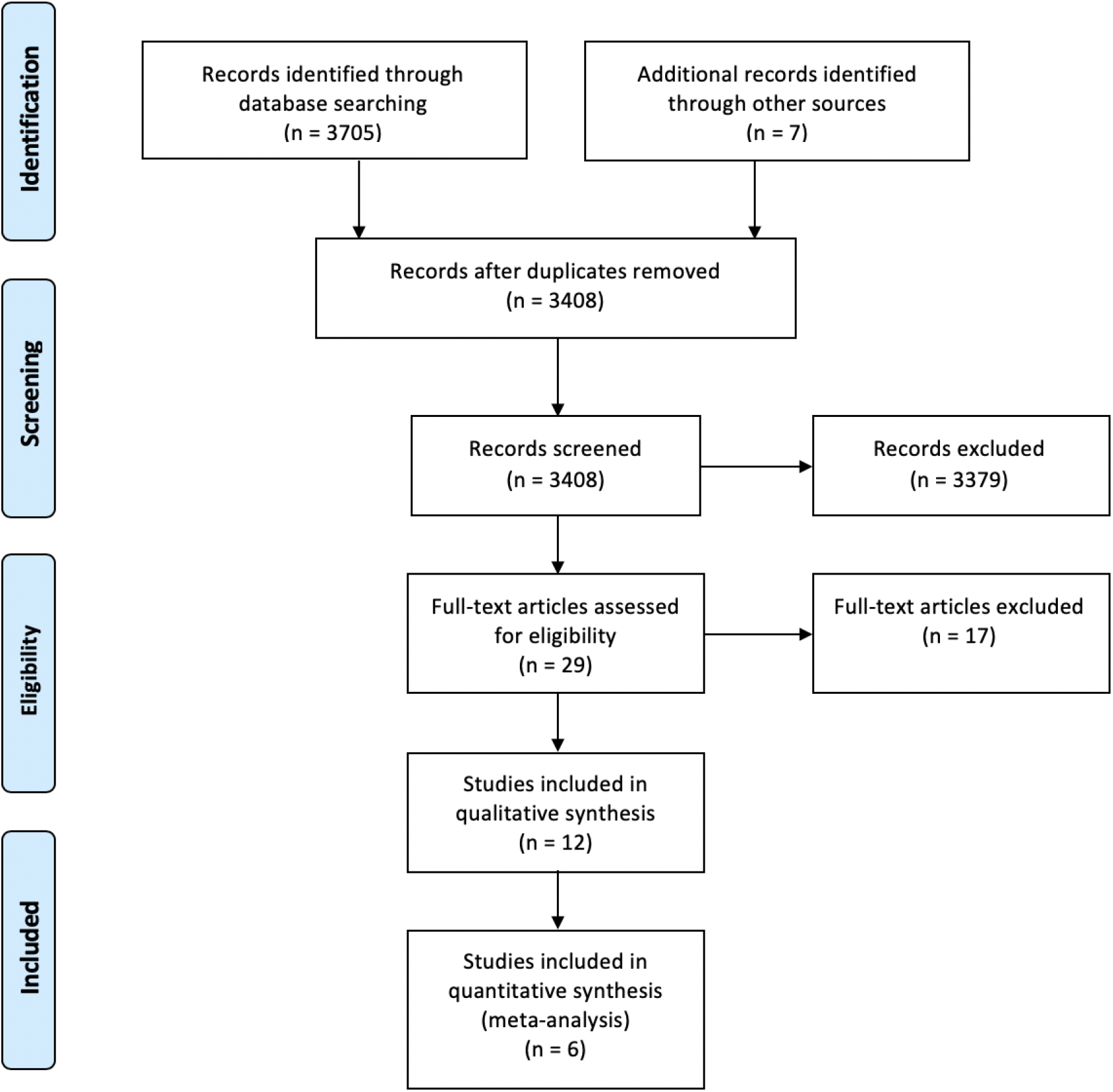
PRISMA flow diagram showing study selection for the systematic literature review and meta-analysis.

We then used a standard data extraction form created in Microsoft Excel, which included fields for information relating to the paper (e.g. authors, year and place of publication), participants and setting (e.g. country, sample size, sex, age, diagnoses), key study details (e.g. type of participants included in the CAH and control group[s], method[s] used for measuring 2D:4D, descriptive statistics for age and digit ratio variables for each participant group [wherever possible]), and a summary of results. When relevant data were missing or ambiguous, the study authors were contacted for clarification. All data were extracted by GR other than for (Nave et al., 2020), which were inputted by SW, and for Turkish language paper (Kocaman et al., 2017), which were extracted by a native Turkish speaker (EA). All data included in the meta-analysis were independently checked by SW, with any disagreements resolved through discussion until a 100% agreement rate was achieved.

### Systematic literature review

Twelve articles examined 2D:4D in CAH populations and were included in the literature review (**Table 1**). Of these, four (Brown et al., 2002; Buck, Williams, Hughes, & Acerini, 2003; Ciumas, Hirschberg, & Savic, 2009; Ökten, Kalyoncu, & Yariş, 2002) were present in the earlier meta-analysis by Hönekopp and Watson (2010), five (Kim et al., 2017; Kocaman et al., 2016, 2017; Oświęcimska et al., 2012; Rivas et al., 2014) had been published since, one (Nave et al., 2020) was currently under review^1^ and two, both relating to the same dataset (Constantinescu et al., 2010; Constantinescu, 2009), were unpublished. All were full-length journal articles other than Kim et al. (2017) and Kocaman et al. (2016), which were published abstracts, and subsequently have appeared as full-length journal articles (Kocaman et al., 2017; Nave et al., 2020), Constantinescu (2009), which was an unpublished MPhil thesis, and Constantinescu et al. (2010), which was a conference poster.

**Table 1.** Studies of 2D:4D in CAH samples included in the systematic literature review.

| Authors | Year | Country | Place of publication | 2D:4D measure(s) | Females with CAH |  | Female controls |  | Males with CAH |  | Male controls |  |
| --- | --- | --- | --- | --- | --- | --- | --- | --- | --- | --- | --- | --- |
|  |  |  |  |  | N | Age | n | Age | n | Age | n | Age |
| Brown et al. | 2002 | UK | *Hormones and Behavior* | Photocopies | 13 | Range = 7–44<br>Average = 15 | 44 | Range = 12–44<br>Average = 18 | 16 | Range = 5–21<br>Average = 11 | 28<sup>1</sup> | Range = 9–34<br>Average = 15 |
| Ökten et al.<sup>2</sup> | 2002 | Turkey | *Early Human Development* | Photocopies, X-rays | 17 | Range = 0–13.3<br>M = 4.6 (SD = 4.2) | 34<sup>3</sup> | Age-matched | 9 | Range = 0–13.3<br>M = 4.6 (SD = 4.2) | 18<sup>3</sup> | Age-matched |
| Buck et al.<sup>4</sup> | 2003 | UK | *Human Reproduction* | X-rays | 66 | Range = 1.1–16.2<br>Median = 8.5 | 69 | Range = 1.9–17<br>Median = 9.3 | 0 | - | 77 | Range = 2.1–20.3<br>Median = 13.86 |
| Ciomas et al. | 2009 | Sweden | *Cerebral Cortex* | Direct (reported)<br>Photocopies (not reported) | 11 | Range = 20–38<br>M = 30 (SD = 8) | 13 | Range = 20–36<br>M = 26 (SD = 7) | 0 | - | 13 | Range = 21–36<br>M = 28 (SD = 6) |
| Constantinescu<sup>a,5</sup> | 2009 | UK | Unpublished MPhil thesis | Direct + photocopies (combined) | 40 | Range = 4–11.83<br>M = 7.50 (SD = 2.20) | 17 | Range = 4.08–12.42<br>M = 7.13 (SD = 2.52) | 24 | Range = 4–11.25<br>M = 7.43 (SD = 1.98) | 7 | Range = 5.08–10.50<br>M = 7.89 (SD = 1.95) |
| Constantinescu et al.<sup>a,6</sup> | 2010 | UK | Unpublished conference poster | Direct + photocopies (combined) | 40 | Range = 4–11.83<br>M = 7.50 (SD = 2.20) | 17 | Range = 4.08–12.42<br>M = 7.13 (SD = 2.52) | 24 | Range = 4–11.25<br>M = 7.43 (SD = 1.98) | 7 | Range = 5.08–10.50<br>M = 7.89 (SD = 1.95) |
| Oświęcimska et al. | 2012 | Poland | *Neuroendocrinology Letters* | X-rays | 19 | Range = 3.7–19<br>M = 13.8 (SD = 4.07) | 0 | - | 0 | - | 0 | - |
| Rivas et al. | 2014 | Brazil | *American Journal of Human Biology* | Direct | 31 | M = 10.7 | 100 | Range = 16–18 | 9 | M = 10.2 | 100 | Range = 16–18 |
| Kocaman et al.<sup>b</sup> | 2016 | Turkey | *Acta Physiologica* | Direct | 30 | Range = 3–15 | 30 | Age-matched | 0 | - | 0 | - |
| Kocaman et al.<sup>b,7</sup> | 2017 | Turkey | *Anadolu Psikiyatri Dergisi* | Direct | 28<sup>8</sup> | M = 8.84 (SD = 4.06)<br>or M = 10.90 (SD = 1.46) | 49 | M = 8.84 (SD = 4.06)<br>or M = 10.90 (SD = 1.46) | 4<sup>8</sup> | M = 8.84 (SD = 4.06)<br>or M = 10.90 (SD = 1.46) | 10 | M = 8.84 (SD = 4.06)<br>or M = 10.90 (SD = 1.46) |
| Kim et al.<sup>c,9</sup> | 2017 | US | *Endocrine Reviews* | X-rays | 40 | Baseline M = 4.6 (SD = 2.8)<br>Follow-up M = 9.9 (SD = 3.3) | 0 | - | 43 | Baseline M = 4.9 (SD = 2.9)<br>Follow-up M = 11.7 (SD = 3.6) | 0 | - |
| Nave et al.<sup>c</sup> | 2020 | US | Under review (*Hormones and Behavior*) | X-rays | 45 | Baseline range: 1.1–18.7 | 31 | Baseline range: 2.6–19.7 | 46 | Baseline range: 1.1–18.7 | 39 | Baseline range: 2.6–19.7 |

Note. <sup>a</sup>, <sup>b</sup>, and <sup>c</sup> indicate sources reporting on the same cohorts.
<sup>1</sup> Although Brown et al.'s (2002) sample includes data from n=28 male controls, L2D:4D was not recorded for n=1 of these participants; therefore, n=27 for L2D:4D, M2D:4D, and D<sub>[R-L]</sub>.
<sup>2</sup> The age-range reported by Ökten et al. (2002) is based on all (male + female) CAH patients; although control participants were matched for age, the exact age-ranges and Ms and SDs were not reported.
<sup>3</sup> Ökten et al. (2002) collected 2D:4D data from n=52 female controls and n=52 male controls but only compared the CAH samples with n=34 age-matched females and n=18 age-matched males.
<sup>4</sup> Buck et al. (2003) examined X-rays for n=71 females with CAH, n=76 female controls, and n=82 male controls (overall, n=17 were rejected because of poor quality films).
<sup>5</sup> Constantinescu (2009) reported that their sample initially consisted of $n=40$ females with CAH (age range = 4-11.83, $M = 7.49$ , $SD = 2.19$ ), $n=25$ males with CAH (age-range = 4-11.25, $M = 7.29$ , $SD = 2.04$ ), $n=18$ female controls (age-range = 4.08-12.42, $M = 7.33$ , $SD = 2.59$ ), and $n=9$ male controls (age-range = 5.08-10.50, $M = 7.62$ , $SD = 1.77$ ). However, 2D:4D data were unavailable for $n=1$ male with CAH, $n=1$ female control, and $n=2$ male controls; we therefore present here the $N$ s, age ranges, $M$ s, and $SD$ s (determined from the original data) based on only those participants for which 2D:4D data were available.
<sup>6</sup> The age range (4-11.8 years) and $M$ age (7.4) reported by Constantinescu et al. (2010) are based on all participants (male + females, with and without CAH); we therefore report here the age ranges, $M$ s and $SD$ s calculated from the original data for each subgroup.
<sup>7</sup> It is unclear what the age of participants was in Kocaman et al. (2017), as the study reports two separate $M$ s and $SD$ s; it is unclear whether these relate to subgroups, and so both $M$ s and $SD$ s are reported here for each group of participants.
<sup>8</sup> Kocaman et al. (2017) reported that they collected data from $n=34$ children with CAH; $n=2$ were removed from analysis because they did not provide consent, and $n=1$ other appears to have been dropped from the analysis (as the overall $n=31$ ), though the reason is unclear.
<sup>9</sup> Age-ranges for Kim et al. (2017) are based on all (male + female) CAH patients.

The studies included in this review were conducted in six countries (Brazil, Poland, Sweden, Turkey, UK, US), and the type of control group to which patients with CAH were compared differed considerably. Some studies employed healthy adult controls without family history of neuropsychiatric conditions (Ciumas et al., 2009), healthy children who had been seen at an outpatient clinic (Ökten et al., 2002), children who had been screened for autism and psychiatric disorders (Kocaman et al., 2016, 2017), otherwise healthy children who had been assessed at an outpatient endocrine clinic due to concerns over short stature (Buck et al., 2003; Nave et al., 2020), university students (Rivas et al., 2014), and unaffected relatives of patients with CAH (Brown et al., 2002; Constantinescu et al., 2010; Constantinescu, 2009; note that some [but not all] of the control participants in Nave et al. [2020] were relatives of their participants with CAH). Some control groups were matched for chronological age (Buck et al., 2003; Ciumas et al., 2009; Ökten et al., 2002) and handedness (Ciumas et al., 2009; Ökten et al., 2002), and one study (Nave et al., 2020) statistically controlled for individual differences in chronological age, bone age, ethnicity, height, puberty status, and ethnicity. For other studies no such controls were implemented (Rivas et al., 2014) or the details are unclear (Kocaman et al., 2016, 2017). Lack of effective control for age between CAH and comparison groups is a point that has been raised as a possible explanation for the inconsistent nature of findings in this literature (McIntyre, Cohn, & Ellison, 2006, p. 149; Nave et al., 2020).

Only one study (Kim et al., 2017) examined whether 2D:4D differs between CAH forms. Although the authors reported no significant difference between patients with classical SW (n=63) or SV (n=20) varieties, further examination is warranted, and particularly so regarding other forms, such as non-classical CAH. Kim et al. (2017) also observed no significant interactions between CAH form, sex, and bone age.

#### Sex differences in CAH samples

Although 2D:4D is lower in males than females in typically developing populations (Hönekopp & Watson, 2010), there was little evidence of such an effect in individual populations with CAH (**Table 2**). No statistically significant effects in the expected (i.e. M<F) direction were observed for either R2D:4D or L2D:4D. A longitudinal study (Kim et al., 2017) also reported no significant sex differences for L2D:4D either at baseline or at final follow-up, or in participants who were pre-pubertal or pubertal. The only study for which significant differences were observed was Rivas et al. (2014), which found the opposite pattern of results than expected (R2D:4D and L2D:4D were both higher in males with CAH than in females with CAH). However, the veracity of these results should be treated with scepticism due to some of the standard deviations reported by Rivas et al. (2014) being much smaller than those of other studies, and the effect size for L2D:4D appearing to be implausibly large (refer to **Table 2** for direct comparison with other studies).

**Table 2.** Comparisons of 2D:4D between males and females with CAH.

| Study | Digit ratio | Measurement | Males |  |  | Females |  |  | Difference |  | *p* | *d* |
| --- | --- | --- | --- | --- | --- | --- | --- | --- | --- | --- | --- | --- |
|  |  |  | *n* | *M* | *SD* | *n* | *M* | *SD* | *t* | *df* |  |  |
| Brown et al. (2002) | R2D:4D<sup>a</sup> | Photocopies | 16 | 0.937 | 0.045 | 13 | 0.957 | 0.038 | -1.235 | 27 | 0.227 | -0.476 |
|  | L2D:4D<sup>a</sup> | Photocopies | 16 | 0.931 | 0.034 | 13 | 0.952 | 0.025 | -1.816 | 27 | 0.080 | -0.692 |
|  | M2D:4D<sup>a</sup> | Photocopies | 16 | 0.934 | 0.037 | 13 | 0.954 | 0.026 | -1.664 | 27 | 0.108 | -0.614 |
|  | D<sub>[R-L]</sub><sup>a</sup> | Photocopies | 16 | 0.006 | 0.033 | 13 | 0.005 | 0.037 | 0.099 | 27 | 0.922 | 0.029 |
| Ökten et al. (2002) | R2D:4D<sup>b</sup> | Photocopies | 9 | 0.92 | 0.04 | 17 | 0.96 | 0.06 | -1.792 | 24 | 0.086 | -0.739 |
|  | L2D:4D<sup>b</sup> | Photocopies | 9 | 0.91 | 0.06 | 17 | 0.92 | 0.05 | -0.453 | 24 | 0.655 | -0.187 |
|  | R2D:4D<sup>c</sup> | X-rays | 9 | 0.98 | 0.03 | 17 | 0.99 | 0.02 | -1.019 | 24 | 0.318 | -0.420 |
|  | L2D:4D<sup>c</sup> | X-rays | 9 | 0.98 | 0.03 | 17 | 0.99 | 0.04 | -0.656 | 24 | 0.518 | -0.271 |
| Constantinescu (2009) | R2D:4D (all subjects)<sup>d,e</sup> | Direct/indirect | 24 | 0.941 | 0.042 | 40 | 0.960 | 0.046 | -1.670 | 62 | 0.100 | -0.426 |
|  | R2D:4D (< 8 years)<sup>d</sup> | Direct/indirect | 15 | 0.933 | 0.042 | 23 | 0.959 | 0.042 | -1.879 | 36 | 0.068 | -0.619 |
|  | R2D:4D (> 8 years)<sup>d,f</sup> | Direct/indirect | 9 | 0.953 | 0.043 | 17 | 0.960 | 0.051 | -0.363 | 24 | 0.720 | -0.144 |
|  | L2D:4D (all subjects)<sup>d,f</sup> | Direct/indirect | 24 | 0.959 | 0.035 | 40 | 0.944 | 0.036 | 1.563 | 62 | 0.123 | 0.421 |
|  | L2D:4D (< 8 years)<sup>d</sup> | Direct/indirect | 15 | 0.958 | 0.040 | 23 | 0.939 | 0.029 | 1.653 | 36 | 0.107 | 0.564 |
|  | L2D:4D (> 8 years)<sup>d,f</sup> | Direct/indirect | 9 | 0.960 | 0.026 | 17 | 0.952 | 0.043 | 0.646 | 23.266<sup>g</sup> | 0.525 | 0.210 |
|  | M2D:4D (all subjects)<sup>d</sup> | Direct/indirect | 24 | 0.950 | 0.028 | 40 | 0.952 | 0.035 | -0.288 | 62 | 0.774 | -0.061 |
|  | M2D:4D (< 8 years)<sup>d</sup> | Direct/indirect | 15 | 0.945 | 0.028 | 23 | 0.949 | 0.031 | -0.392 | 36 | 0.697 | -0.134 |
|  | M2D:4D (> 8 years)<sup>d</sup> | Direct/indirect | 9 | 0.957 | 0.030 | 17 | 0.956 | 0.042 | 0.048 | 24 | 0.962 | 0.026 |
|  | D<sub>[R-L]</sub> (all subjects)<sup>d</sup> | Direct/indirect | 24 | -0.018 | 0.053 | 40 | 0.016 | 0.041 | -2.811 | 62 | **0.007** | **-0.742** |
|  | D<sub>[R-L]</sub> (< 8 years)<sup>d</sup> | Direct/indirect | 15 | -0.025 | 0.061 | 23 | 0.020 | 0.039 | -2.781 | 36 | **0.009** | **-0.923** |
|  | D<sub>[R-L]</sub> (> 8 years)<sup>d</sup> | Direct/indirect | 9 | -0.007 | 0.038 | 17 | 0.009 | 0.045 | -0.910 | 24 | 0.372 | -0.374 |
| Rivas et al. (2014) | R2D:4D<sup>g</sup> | Direct | 9 | 0.960 | 0.0220 | 31 | 0.950 | 0.0077 | 2.166 | 38 | **0.037** | **0.820** |
|  | L2D:4D<sup>g</sup> | Direct | 9 | 0.983 | 0.0267 | 31 | 0.947 | 0.0114 | 5.981 | 38 | **< 0.001** | **2.265** |
| Kim et al. (2017) | L2D:4D (baseline) | X-rays | 43 | 0.902 | 0.035 | 40 | 0.911 | 0.026 | -1.260 | 81 | 0.211 | - |
|  | L2D:4D (follow-up) | X-rays | 43 | 0.918 | 0.026 | 40 | 0.920 | 0.025 | -0.396 | 81 | 0.693 | - |
|  | L2D:4D (pre-pubertal) | X-rays | 16 | 0.89 | 0.04 | 19 | 0.91 | 0.02 | 1.917 | 33 | 0.064 | - |
|  | L2D:4D (pubertal) | X-rays | 27 | 0.92 | 0.03 | 15 | 0.92 | 0.03 | 0.0 | 40 | 1.0 | - |
| Nave et al. (2020) | L2D:4D (average) | X-rays | 45 | 0.913 | 0.023 | 45 | 0.917 | 0.023 | -0.949 | 88 | 0.345 | -0.200 |
|  | L2D:4D (first scan) | X-rays | 45 | 0.902 | 0.033 | 45 | 0.911 | 0.027 | -1.380 | 88 | 0.171 | -0.291 |
|  | L2D:4D (last scan) | X-rays | 45 | 0.918 | 0.024 | 45 | 0.920 | 0.023 | -0.550 | 88 | 0.584 | -0.116 |

Note. Negative *d* values indicate effects in the predicted direction (i.e. $M < F$ ); effects in bold are statistically significant ( $p < 0.05$ ).
<sup>a</sup> = calculated from original data of Brown et al. (2002)
<sup>b</sup> = calculated from data presented by Ökten et al. (2002, Table 1, p. 50)
<sup>c</sup> = calculated from data presented by Ökten et al. (2002, Table 3, p. 51)
<sup>d</sup> = calculated from the original data of Constantinescu (2009)
<sup>e</sup> = we report here the values calculated from the original data due to there being differences in rounding with the values reported by Constantinescu (2009); the values reported by Constantinescu are as follows: males with CAH ( $n=24$ , $M = 0.9407$ , $SD = 0.04$ ), females with CAH ( $n=40$ , $M = 0.9598$ , $SD = 0.04$ ), $t(62) = -1.67$ , $p = 0.100$ , $d = -0.43$
<sup>f</sup> = we report here the values calculated from the original data due to there being differences in rounding with the values reported by Constantinescu (2009); the values reported by Constantinescu are as follows: males with CAH ( $n=24$ , $M = 0.9586$ , $SD = 0.03$ ), females with CAH ( $n=40$ , $M = 0.9444$ , $SD = 0.03$ ), $t(62) = 1.56$ , $p = 0.123$ , $d = 0.40$ .
<sup>g</sup> = calculated from data presented by Rivas et al. (2014, p. 560)
<sup>†</sup> = in Constantinescu (2009), it is listed that the $> 8$ years old group consisted of 19 females and 7 males, whereas in the dataset we analysed, there were 17 females and 9 males.
<sup>§</sup> = equal variances not assumed

No studies reported whether M2D:4D (i.e. the average of R2D:4D and L2D:4D) or D_[R-L]_ (i.e. the right-left difference in 2D:4D) differed between males with CAH and females with CAH. Reanalysis of the datasets presented by Brown et al. (2002) and Constantinescu (2009) yielded no significant sex differences for M2D:4D. Although we also found no sex difference for D_[R-L]_ in Brown et al.’s (2002) data, this was not the case for Constantinescu (2009): D_[R-L]_ was significantly lower in males with CAH than in females with CAH. This appeared to be driven by a significant difference in the younger group (< 8 years old), as no such effect was detected in the older group (> 8 years old), and could potentially therefore reflect differences in bone maturation rates.

#### Other reported correlations within CAH samples

**Table 3** presents additional findings from studies of 2D:4D in CAH samples. Two studies (Buck et al., 2003; Kim et al., 2017; Nave et al., 2020) reported associations between 2D:4D and age in patients with CAH. Of note, the only longitudinal study in the area (Kim et al., 2017; Nave et al., 2020) reported that L2D:4D increased between baseline and final follow-up, and that the effect size (*d* = 0.46) was small-medium (Cohen, 1988; 0.20 = small, 0.50 = medium, 0.80 = large); further, 2D:4D was lower in pre-pubertal than pubertal participants. Buck et al. (2003) also reported that 2D:4D correlated positively with age in their cohort, though noted that the effect was not statistically significant. Positive correlations between 2D:4D and age are consistent with the findings of some cross-sectional (e.g. Richards, Bellin, & Davies, 2017) and longitudinal studies (McIntyre, Ellison, Lieberman, Demerath, & Towne, 2005; Trivers, Manning, & Jacobson, 2006) of typically developing children.

**Table 3.** Additional findings from studies of 2D:4D in CAH populations.

| Authors | Main findings |
| --- | --- |
| Buck et al. (2003) | L2D:4D marginally increased with age ($p = 0.08$) (analysis included females with CAH, and male and female controls) |
| Constantinescu (2009) | No significant correlation between R2D:4D of males with CAH and R2D:4D of their mothers<br>No significant correlation between L2D:4D of males with CAH and L2D:4D of their mothers<br>R2D:4D in females with CAH correlated positively with R2D:4D in their mothers<br>R2D:4D in females with CAH correlated positively with L2D:4D in their mothers<br>No significant correlation between L2D:4D of females with CAH and L2D:4D of their mothers<br>L2D:4D in females with CAH correlated positively with R2D:4D in their mothers<br>†No significant correlation between age and R2D:4D in males<br>†No significant correlation between age and L2D:4D in males<br>†No significant correlation between age and M2D:4D in males<br>†No significant correlation between age and $D_{[R-L]}$ in males<br>†No significant correlation between age and R2D:4D in females<br>†No significant correlation between age and L2D:4D in females<br>†No significant correlation between age and M2D:4D in females<br>†Significant negative correlation between age and $D_{[R-L]}$ in females |
| Oświęcimska et al. (2012) | Mean 2D:4D in females with CAH correlated positively with serum testosterone<br>Mean 2D:4D in females with CAH correlated positively with serum s-DHEA<br>Mean 2D:4D in females with CAH did not correlate with serum androstenedione |
| Kocaman et al. (2017) | 2D:4D correlated with autistic traits (direction unclear; hand unclear)<br>2D:4D in children with CAH was related to maternal stress (direction unclear; hand unclear)<br>2D:4D did not differ in children who had a difficult birth (hand unclear)<br>2D:4D did not differ in children who had a premature birth (hand unclear)<br>2D:4D did not differ in children whose mothers smoked (hand unclear)<br>2D:4D did not differ in children whose mothers had a physical or mental health condition (hand unclear) |
| Nave et al. (2020) | L2D:4D lower in Hispanic than White participants<br>L2D:4D correlated positively with bone age in males<br>L2D:4D correlated positively with bone age in females<br>L2D:4D correlated negatively with puberty stage<br>L2D:4D correlated positively with height<br>No interaction between sex and CAH |

*Note.* Kocaman et al. (2016) is not included in this Table because it was an earlier publication of the same study presented by Kocaman et al. (2017); Kim et al., (2017) is also not included because it was an earlier publication of data from Nave et al. (2020); Constantinescu et al. (2010) is not included because it is a less complete report of the study by Constantinescu (2009).
† Effect not reported in the original article (Pearson's correlations were calculated from the original dataset of Constantinescu [2009]).

Although Constantinescu (2009) reported subgroup analyses based on age, they did not report whether 2D:4D correlated with age. We therefore conducted Pearson’s correlations to examine this possibility. In males, age did not correlate significantly with R2D:4D (*r*[22] = 0.178, *p* = 0.405), L2D:4D (*r*[22] = −0.275, *p* = 0.194), M2D:4D (*r*[22] = −0.036, *p* = 0.868), or D_[R-L]_ (*r*[22] = 0.323, *p* = 0.124). In females, age correlated negatively with D_[R-L]_ (*r*[38] = −0.320, *p* = 0.044, though did not correlate significantly with R2D:4D (*r*[38] = −0.174, *p* = 0.283), L2D:4D (*r*[38] = 0.149, *p* = 0.359), or M2D:4D (*r*[38] = −0.037, *p* = 0.819).

Unlike other studies, Kocaman et al. (2016, 2017) presented analyses in which the male and female samples were combined. These authors reported that a Pearson’s correlation between 2D:4D and a measure of autistic traits (a Turkish language translation of the Autism Behavior Checklist [ABC]) was significant (direction of effect is unclear). It is also ambiguous whether this effect was observed in CAH participants or controls, and whether it related to R2D:4D or L2D:4D (the effect examined in the analysis in which patients with CAH and controls were combined was not significant). Although the finding is difficult to interpret, it may be relevant in regard to previous studies that have reported correlations between 2D:4D, autism, and autistic traits (Hönekopp, 2012; Manning, Baron-Cohen, Wheelwright, & Sanders, 2001; Myers, van’t Westeinde, Kuja-Halkola, Tammimies, & Bölte, 2018; Schieve et al., 2018; Teatero & Netley, 2013; Voracek, 2008). Kocaman et al. (2017) also reported that 2D:4D did not differ for children who had a difficult birth, a premature birth, or whose mother smoked or had a physical or mental health condition. There was, however, a significant effect of maternal stress within the CAH group, though the direction of this effect is unclear.

Although endocrine status is frequently monitored in patients with CAH, the only study so far to report on circulating hormone levels and 2D:4D in a CAH sample is Oświęcimska et al. (2012). These authors reported that M2D:4D was positively correlated with serum testosterone and dehydroepiandrostenedione sulphate (DHEAS), though there was no association with androstenedione, and they did not report whether there was a correlation with 17-hydroxyprogesterone (17-OHP). (Also note that although both the Abstract and Results sections of this paper report that M2D:4D correlated positively with testosterone and s-DHEA, Figures 1 and 2 reportedly present significant positive correlations between M2D:4D and testosterone and androstenedione, respectively.) These findings are difficult to interpret, as they relate to a small sample, and no other published study has examined such effects in a CAH population. Although some individual studies have reported significant correlations between 2D:4D and circulating testosterone, meta-analyses suggest these variables are not related (Hönekopp, Bartholdt, Beier, & Liebert, 2007; Zhang et al., 2019). It is therefore suggested that unless these effects are replicated, they should be interpreted with caution.

**Figure 2.**
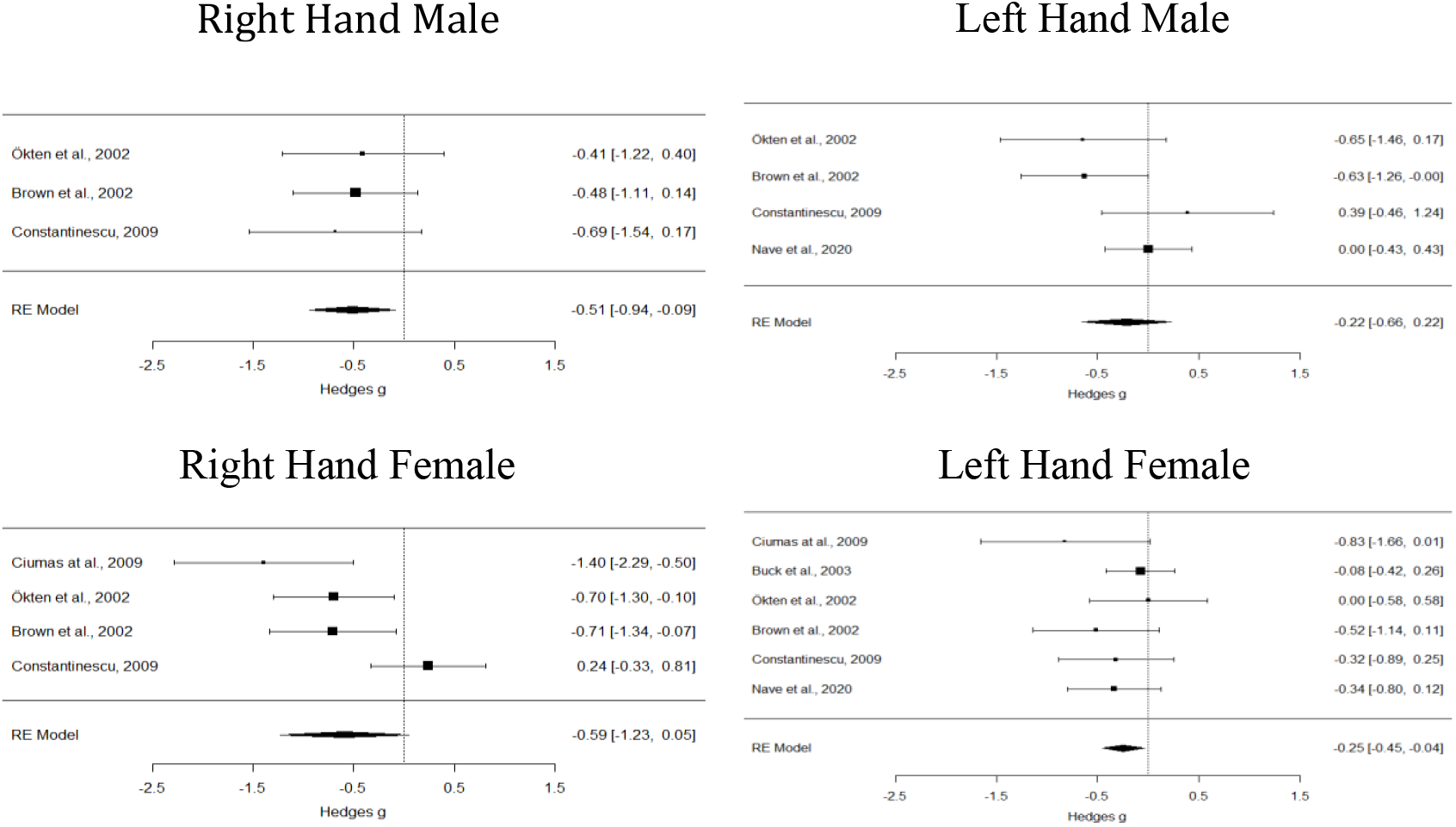
Forest plot summary for each meta-analysis comparing 2D:4D for individuals with CAH to controls for each sex and hand combination.

#### Comparisons of 2D:4D between patients with CAH and controls

Findings from studies comparing 2D:4D between CAH samples and control samples are reported in **Table 4**. Significantly lower 2D:4D has been reported in five CAH cohorts (Brown et al., 2002; Ciumas et al., 2009; Kocaman et al., 2016, 2017; Ökten et al., 2002; Rivas et al., 2014). However, three studies (Buck et al., 2003; Constantinescu et al., 2010; Constantinescu, 2009; Nave et al., 2020) reported only null-findings. Notably, these included the largest (Buck et al., 2003: n=66), second largest (Nave et al., 2020; n=45), and third largest (Constantinescu, 2009; n=40) samples of females with CAH, as well as the largest (Nave et al., 2020; n=45) and second largest (Constantinescu, 2009; n=24) samples of males with CAH. However, to interpret these findings accurately, some further consideration of the studies’ methodologies is required. Constantinescu and colleagues used an unusual approach for measuring 2D:4D: a combination of both direct (calliper) and indirect (photocopy) measures were collected, with both types of measurements being recorded for a subset of participants. For those from whom only direct measures were available, these were then adjusted so that they resembled photocopy measures. They did this by dividing the overall mean value from the photocopy measurements by the mean for the calliper measurements, then multiplying this by the calliper measurement for each individual. Additionally, although the CAH samples were relatively large (female n=40, male n=24), the comparison samples were not (female n=17, male n=7), meaning that the benefit in terms of statistical power associated with large CAH samples was somewhat undermined by the small control groups. Buck et al. (2003) on the other hand did not examine males with CAH, only recorded L2D:4D (and not R2D:4D), and measured digit ratios from X-rays. Likewise, Nave et al. (2020) compared only L2D:4D (from X-rays) between patients with CAH and controls (although they did examine both males and females).

**Table 4.**
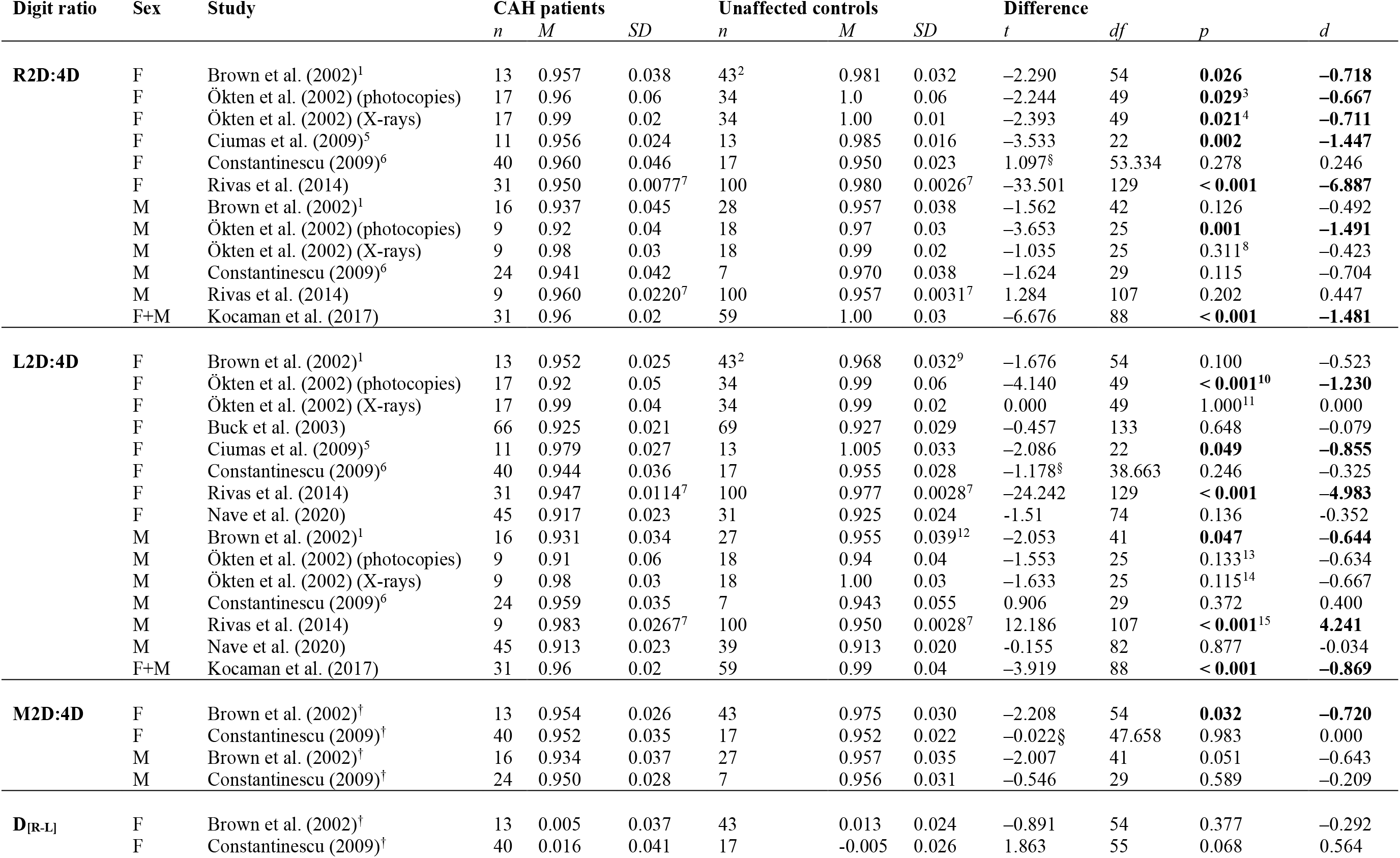
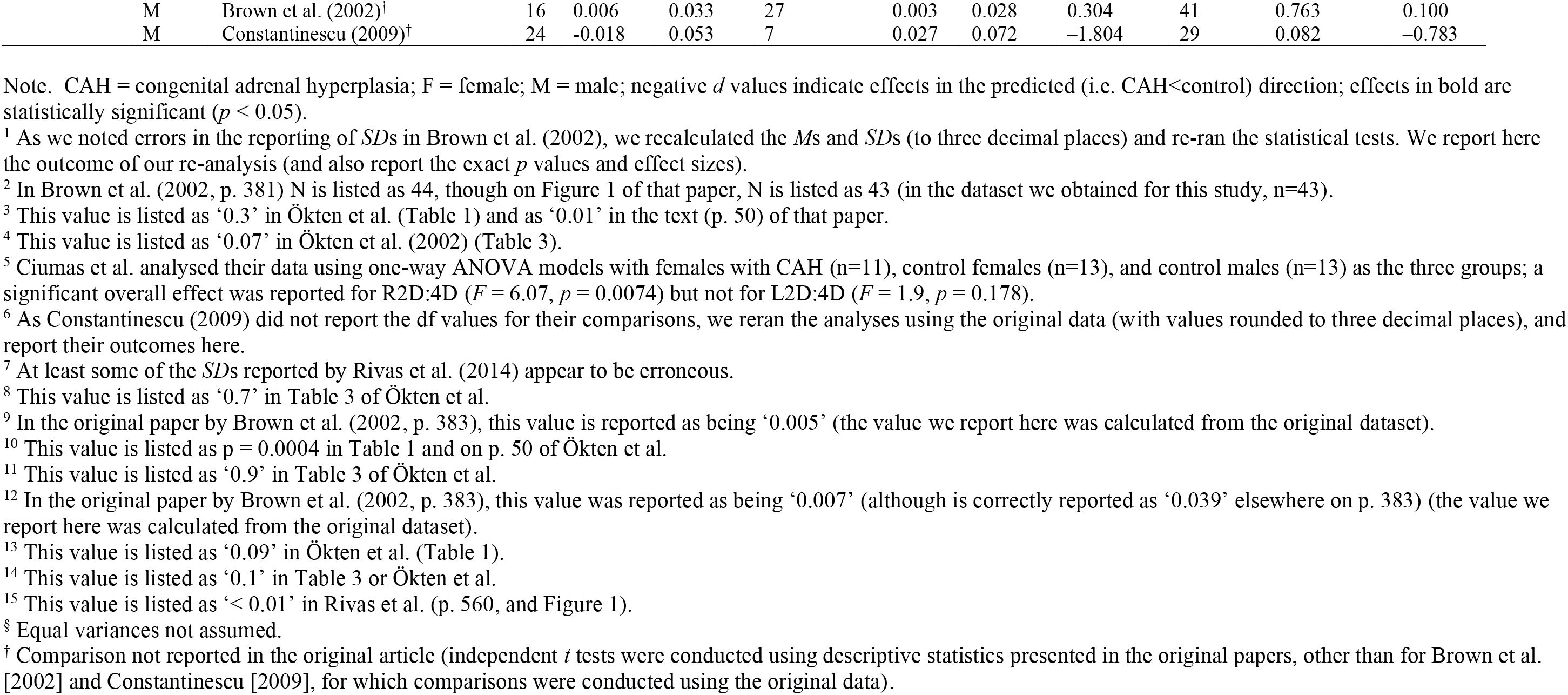
Comparisons of digit ratio variables between patients with CAH and unaffected controls.

| Digit ratio | Sex | Study | CAH patients |  |  | Unaffected controls |  |  | Difference |  |  |  |
| --- | --- | --- | --- | --- | --- | --- | --- | --- | --- | --- | --- | --- |
|  |  |  | *n* | *M* | *SD* | *n* | *M* | *SD* | *t* | *df* | *p* | *d* |
| **R2D:4D** | F | Brown et al. (2002)<sup>1</sup> | 13 | 0.957 | 0.038 | 43<sup>2</sup> | 0.981 | 0.032 | -2.290 | 54 | **0.026** | **-0.718** |
|  | F | Ökten et al. (2002) (photocopies) | 17 | 0.96 | 0.06 | 34 | 1.0 | 0.06 | -2.244 | 49 | **0.029<sup>3</sup>** | **-0.667** |
|  | F | Ökten et al. (2002) (X-rays) | 17 | 0.99 | 0.02 | 34 | 1.00 | 0.01 | -2.393 | 49 | **0.021<sup>4</sup>** | **-0.711** |
|  | F | Ciomas et al. (2009)<sup>5</sup> | 11 | 0.956 | 0.024 | 13 | 0.985 | 0.016 | -3.533 | 22 | **0.002** | **-1.447** |
|  | F | Constantinescu (2009)<sup>6</sup> | 40 | 0.960 | 0.046 | 17 | 0.950 | 0.023 | 1.097<sup>§</sup> | 53.334 | 0.278 | 0.246 |
|  | F | Rivas et al. (2014) | 31 | 0.950 | 0.0077<sup>7</sup> | 100 | 0.980 | 0.0026<sup>7</sup> | -33.501 | 129 | **< 0.001** | **-6.887** |
|  | M | Brown et al. (2002)<sup>1</sup> | 16 | 0.937 | 0.045 | 28 | 0.957 | 0.038 | -1.562 | 42 | 0.126 | -0.492 |
|  | M | Ökten et al. (2002) (photocopies) | 9 | 0.92 | 0.04 | 18 | 0.97 | 0.03 | -3.653 | 25 | **0.001** | **-1.491** |
|  | M | Ökten et al. (2002) (X-rays) | 9 | 0.98 | 0.03 | 18 | 0.99 | 0.02 | -1.035 | 25 | 0.311<sup>8</sup> | -0.423 |
|  | M | Constantinescu (2009)<sup>6</sup> | 24 | 0.941 | 0.042 | 7 | 0.970 | 0.038 | -1.624 | 29 | 0.115 | -0.704 |
|  | M | Rivas et al. (2014) | 9 | 0.960 | 0.0220<sup>7</sup> | 100 | 0.957 | 0.0031<sup>7</sup> | 1.284 | 107 | 0.202 | 0.447 |
|  | F+M | Kocaman et al. (2017) | 31 | 0.96 | 0.02 | 59 | 1.00 | 0.03 | -6.676 | 88 | **< 0.001** | **-1.481** |
| **L2D:4D** | F | Brown et al. (2002)<sup>1</sup> | 13 | 0.952 | 0.025 | 43<sup>2</sup> | 0.968 | 0.032<sup>9</sup> | -1.676 | 54 | 0.100 | -0.523 |
|  | F | Ökten et al. (2002) (photocopies) | 17 | 0.92 | 0.05 | 34 | 0.99 | 0.06 | -4.140 | 49 | **< 0.001<sup>10</sup>** | **-1.230** |
|  | F | Ökten et al. (2002) (X-rays) | 17 | 0.99 | 0.04 | 34 | 0.99 | 0.02 | 0.000 | 49 | 1.000<sup>11</sup> | 0.000 |
|  | F | Buck et al. (2003) | 66 | 0.925 | 0.021 | 69 | 0.927 | 0.029 | -0.457 | 133 | 0.648 | -0.079 |
|  | F | Ciomas et al. (2009)<sup>5</sup> | 11 | 0.979 | 0.027 | 13 | 1.005 | 0.033 | -2.086 | 22 | **0.049** | **-0.855** |
|  | F | Constantinescu (2009)<sup>6</sup> | 40 | 0.944 | 0.036 | 17 | 0.955 | 0.028 | -1.178<sup>§</sup> | 38.663 | 0.246 | -0.325 |
|  | F | Rivas et al. (2014) | 31 | 0.947 | 0.0114<sup>7</sup> | 100 | 0.977 | 0.0028<sup>7</sup> | -24.242 | 129 | **< 0.001** | **-4.983** |
|  | F | Nave et al. (2020) | 45 | 0.917 | 0.023 | 31 | 0.925 | 0.024 | -1.51 | 74 | 0.136 | -0.352 |
|  | M | Brown et al. (2002)<sup>1</sup> | 16 | 0.931 | 0.034 | 27 | 0.955 | 0.039<sup>12</sup> | -2.053 | 41 | **0.047** | **-0.644** |
|  | M | Ökten et al. (2002) (photocopies) | 9 | 0.91 | 0.06 | 18 | 0.94 | 0.04 | -1.553 | 25 | 0.133<sup>13</sup> | -0.634 |
|  | M | Ökten et al. (2002) (X-rays) | 9 | 0.98 | 0.03 | 18 | 1.00 | 0.03 | -1.633 | 25 | 0.115<sup>14</sup> | -0.667 |
|  | M | Constantinescu (2009)<sup>6</sup> | 24 | 0.959 | 0.035 | 7 | 0.943 | 0.055 | 0.906 | 29 | 0.372 | 0.400 |
|  | M | Rivas et al. (2014) | 9 | 0.983 | 0.0267<sup>7</sup> | 100 | 0.950 | 0.0028<sup>7</sup> | 12.186 | 107 | **< 0.001<sup>15</sup>** | **4.241** |
|  | M | Nave et al. (2020) | 45 | 0.913 | 0.023 | 39 | 0.913 | 0.020 | -0.155 | 82 | 0.877 | -0.034 |
|  | F+M | Kocaman et al. (2017) | 31 | 0.96 | 0.02 | 59 | 0.99 | 0.04 | -3.919 | 88 | **< 0.001** | **-0.869** |
| **M2D:4D** | F | Brown et al. (2002)<sup>†</sup> | 13 | 0.954 | 0.026 | 43 | 0.975 | 0.030 | -2.208 | 54 | **0.032** | **-0.720** |
|  | F | Constantinescu (2009)<sup>†</sup> | 40 | 0.952 | 0.035 | 17 | 0.952 | 0.022 | -0.022<sup>§</sup> | 47.658 | 0.983 | 0.000 |
|  | M | Brown et al. (2002)<sup>†</sup> | 16 | 0.934 | 0.037 | 27 | 0.957 | 0.035 | -2.007 | 41 | 0.051 | -0.643 |
|  | M | Constantinescu (2009)<sup>†</sup> | 24 | 0.950 | 0.028 | 7 | 0.956 | 0.031 | -0.546 | 29 | 0.589 | -0.209 |
| **D<sub>[R-L]</sub>** | F | Brown et al. (2002)<sup>†</sup> | 13 | 0.005 | 0.037 | 43 | 0.013 | 0.024 | -0.891 | 54 | 0.377 | -0.292 |
|  | F | Constantinescu (2009)<sup>†</sup> | 40 | 0.016 | 0.041 | 17 | -0.005 | 0.026 | 1.863 | 55 | 0.068 | 0.564 |

|  |  |  |  |  |  |  |  |  |  |  |  |
| --- | --- | --- | --- | --- | --- | --- | --- | --- | --- | --- | --- |
| M | Brown et al. (2002)<sup>†</sup> | 16 | 0.006 | 0.033 | 27 | 0.003 | 0.028 | 0.304 | 41 | 0.763 | 0.100 |
| M | Constantinescu (2009)<sup>†</sup> | 24 | -0.018 | 0.053 | 7 | 0.027 | 0.072 | -1.804 | 29 | 0.082 | -0.783 |

Note. CAH = congenital adrenal hyperplasia; F = female; M = male; negative *d* values indicate effects in the predicted (i.e. CAH<control) direction; effects in bold are statistically significant ( $p < 0.05$ ).
<sup>1</sup> As we noted errors in the reporting of *SDs* in Brown et al. (2002), we recalculated the *Ms* and *SDs* (to three decimal places) and re-ran the statistical tests. We report here the outcome of our re-analysis (and also report the exact *p* values and effect sizes).
<sup>2</sup> In Brown et al. (2002, p. 381) *N* is listed as 44, though on Figure 1 of that paper, *N* is listed as 43 (in the dataset we obtained for this study, *n*=43).
<sup>3</sup> This value is listed as ‘0.3’ in Ökten et al. (Table 1) and as ‘0.01’ in the text (p. 50) of that paper.
<sup>4</sup> This value is listed as ‘0.07’ in Ökten et al. (2002) (Table 3).
<sup>5</sup> Ciumas et al. analysed their data using one-way ANOVA models with females with CAH (*n*=11), control females (*n*=13), and control males (*n*=13) as the three groups; a significant overall effect was reported for R2D:4D ( $F = 6.07$ , $p = 0.0074$ ) but not for L2D:4D ( $F = 1.9$ , $p = 0.178$ ).
<sup>6</sup> As Constantinescu (2009) did not report the *df* values for their comparisons, we reran the analyses using the original data (with values rounded to three decimal places), and report their outcomes here.
<sup>7</sup> At least some of the *SDs* reported by Rivas et al. (2014) appear to be erroneous.
<sup>8</sup> This value is listed as ‘0.7’ in Table 3 of Ökten et al.

Brown et al. (2002) provided evidence to suggest the difference in 2D:4D between patients with CAH and controls may be due to environmental influences (e.g. the elevated prenatal testosterone exposure characteristic of CAH) rather than shared genetics, as they observed lower 2D:4D in male patients with CAH than in their unaffected male relatives (R2D:4D, *p* = 0.033, *d* = −1.191; L2D:4D, *p* = 0.011, *d* = − 1.592) (Note that these effects are incorrectly reported in the original paper as R2D:4D, *p* = 0.01, *d* = 1.0; L2D:4D, *p* < 0.04, *d* = 0.9). However, this analysis relied on a very small sample (males with CAH, n=6; unaffected males, n=6), and a larger study (Constantinescu et al., 2010; Constantinescu, 2009) found no such differences in males or females (males with CAH, n=24; unaffected males, n=9; females with CAH, n=40; unaffected females, n=18). Interestingly though, Constantinescu (2009) reported only weak to moderate sized correlations (many of which were not statistically significant) between digit ratios of children and those of their mothers, whereas previous studies (e.g. Hiraishi, Sasaki, Shikishima, & Ando, 2012; Kalichman, Batsevich, & Kobyliansky, 2019; Voracek & Dressler, 2009) suggest that genetic factors contribute substantially to the phenotypic expression of this trait.

The only study to observe a significant effect in the opposite direction than expected was Rivas et al. (2014), who reported L2D:4D in males with CAH to be higher than that of male controls. However, only 9 males with CAH were included in this analysis (in comparison to 100 male controls), and the mean age of the CAH males was 10.2 years whereas the control group consisted of students aged 16-18 years. This sample was also reported to be ethnically diverse, which could be important considering that 2D:4D can vary more by ethnicity than by sex (de Sanctis et al., 2017; Loehlin, McFadden, Medland, & Martin, 2006; Manning, Churchill, & Peters, 2007; Manning, Stewart, Bundred, & Trivers, 2004). In addition, and likely of greater importance, there appear to be errors in the reporting of some of the standard deviations (i.e. some were implausibly small) (see text on p. 560 as well as the error bars on Figure 1 of that paper). This observation renders these results unreliable and therefore they are not interpreted further.

Ökten et al. (2002) reported that 10 girls with CAH who were less than 2 years old had significantly lower R2D:4D and L2D:4D than age-matched female controls. This could suggest that differences in 2D:4D appear early in life, which is consistent with the idea that they relate to prenatal androgen exposure. However, Constantinescu (2009) found only marginally (*p* = 0.063, *d* = 0.72) lower L2D:4D in girls with CAH compared to their unaffected female relatives aged 4–7.99 years, and no difference for R2D:4D; there were also no differences for R2D:4D or L2D:4D in boys of this age. No differences were observed between girls with CAH and unaffected girls or between boys with CAH and unaffected boys aged 8–12.42 years for either R2D:4D or L2D:4D.

No studies reported whether M2D:4D or D_[R-L]_ differed between CAH populations and controls. However, we were able to conduct such analyses from the original data of Brown et al. (2002) and Constantinescu (2009) (see **Table 4**). For Brown et al. (2002), we observed that M2D:4D was significantly lower in females with CAH than female controls. M2D:4D was also lower in males with CAH than male controls, though the effect was just short of statistical significance (*p* = 0.051, *d* = 0.633). A paired-samples *t* test determined that M2D:4D was significantly lower in males with CAH (n=6, *M* = 0.911, *SD* = 0.042) than in their unaffected male relatives (n=6, *M* = 0.955, *SD* = 0.033), *t*(5) = −4.043, *p* = 0.01, *d* = 1.164. However, in Constantinescu’s (2009) data, there was no difference in M2D:4D between females with CAH and unaffected females; there was also no difference between males with CAH and unaffected males.

When examining D_[R-L]_ in Brown et al.’s (2002) dataset, we found no significant differences between females with CAH and female controls, or between males with CAH and male controls. A paired-samples *t* test determined that D_[R-L]_ also did not differ between males with CAH (n=6, *M* = −0.008, *SD* = 0.032) and their unaffected male relatives (n=6, *M* = 0.004, *SD* = 0.041), *t*(5) = −0.704, *p* = 0.513, *d* = 0.336. In Constantinescu’s (2009) dataset, D_[R-L]_ was marginally lower in males with CAH than unaffected males (*p* = 0.082). However, marginally higher D_[R-L]_ was observed in females with CAH compared to unaffected females (*p* = 0.068).

### Meta-analysis

To determine whether R2D:4D and L2D:4D differ significantly between females with CAH and control females, and between males with CAH and control males, we conducted meta-analyses using the R package metafor (Viechtbauer, 2010).

The inclusion criteria for the meta-analysis were that studies had to report primary 2D:4D data for humans with CAH as well as for controls, and that they needed to report effect sizes and/or statistics from which effect sizes could be calculated. We contacted the first/corresponding authors of the relevant studies to request the data necessary to calculate effect sizes if they were not available within the published articles. If we did not hear back, we subsequently contacted other authors for whom contact details could be obtained. The standard deviations reported in Brown et al. (2002) were clearly standard errors (and we checked this using the original dataset), so we were able to calculate the correct values and include this study. Rivas et al. (2014) also reported standard deviations that very likely contain an error given the implausibly large effect sizes generated in this study, and standard deviations far smaller for some subsamples than is typical in 2D:4D literature. However, unlike with Brown et al. (2002), it was not obvious that the reported values definitely referred to standard errors and so we excluded this study from the meta-analysis.

Random-effects models with the restricted maximum-likelihood estimation method were calculated to account for likely heterogeneity in the data. To best compensate for our small samples, we report standardised mean differences in the form of Hedges’ *g* (Hedges & Olkin, 1985). We report heterogeneity in terms of I^2^. For completeness, we also report Cochran’s Q as a formal test for the presence of heterogeneity, though caution that this test is likely underpowered due to the low number of relevant studies identified.

Egger’s regression was used to formally test the potential for small study effects and publication bias and we illustrate these using funnel plots. Only two studies provided estimates for 2D:4D that aggregate across both hands, so it was not possible to perform Egger’s regression for these estimates. Due to the low number of studies we did not perform any formal tests of potential sources of heterogeneity via subgroup analyses or meta-regression. Instead we refer the reader to the systematic review where possible sources of heterogeneity are described in detail (Section 3).

#### CAH vs. Control Participants Meta-Analysis Results

We present the results of meta-analyses comparing differences in individual hands via Forest Plots in **Figure 2**, and for aggregated measures (M2D:4D and D_[R-L_]) in **Figure 3**. We present summary statistics for all meta-analyses in **Table 5**. Only two comparisons identified statistically significant differences between individuals with and without CAH: the right-hand comparison for males, and the left-hand comparison for females. Egger’s test of small study effects did not identify statistically significant effects for any analysis. However, the small number of studies provided low power for this test. We present funnel plots in **Figure 4**, some of which do suggest that small study effects were plausible for some comparisons. For female samples the right-hand analysis was not far from statistical significance (*z* = −1.877, *p* = 0.061), while there were no indications of small study effects for the left hand (*z* = −1.552, *p* = 0.121). Male samples did not show any sign of small study effects for right or left hand (*z* = −0.206, *p* = 0.837, and *z* = −0.067, *p* = 0.947, respectively). Notably, the effect sizes observed in these meta-analyses were considerably smaller than those reported by Hönekopp and Watson (2010) (for comparisons, see **Table 6**). We therefore conducted a priori power analyses using G*Power 3.1 (Faul, Erdfelder, Lang, & Buchner, 2007) based on the effect sizes observed in the present study (note that we substituted *d* for *g* in this case, the difference being negligible) to determine the number of participants that would be required to observe statistically significant (*p* < 0.05, two-tailed) effects with 80% power using independent samples t-tests with equal numbers of cases and controls. The required sample sizes for R2D:4D (males: n=122; females: n=92) were considerably smaller than those for L2D:4D (males: n=664; females, n=526).

**Table 5.** Summary of meta analyses of the difference between 2D:4D for participants with CAH and controls.

| Digit Ratio | Sex | Study | CAH |  |  | Control |  |  | Effect Size |  | Meta-Analyses (95%CI) |  |  |  | Heterogeneity |  |  |  |  |  |
| --- | --- | --- | --- | --- | --- | --- | --- | --- | --- | --- | --- | --- | --- | --- | --- | --- | --- | --- | --- | --- |
|  |  |  | *n* | *M* | *SD* | *n* | *M* | *SD* | *g* | *SE* | *g* | LCI | UCI | *SE* | *p* | Q | df | *p* | τ | I<sup>2</sup> |
| R2D:4D | Male | Ökten et al. | 9 | 0.980 | 0.030 | 18 | 0.990 | 0.020 | -0.410 | 0.412 | -0.513 | -0.940 | -0.085 | 0.218 | 0.019 | 0.226 | 2 | 0.893 | 0 | 0 |
|  |  | Brown et al. | 16 | 0.937 | 0.045 | 28 | 0.957 | 0.038 | -0.483 | 0.318 |  |  |  |  |  |  |  |  |  |  |
|  |  | Constantinescu et al. | 24 | 0.941 | 0.042 | 7 | 0.970 | 0.038 | -0.686 | 0.439 |  |  |  |  |  |  |  |  |  |  |
|  | Female | Ciumas et al | 11 | 0.956 | 0.024 | 13 | 0.985 | 0.016 | -1.398 | 0.460 | -0.591 | -1.233 | 0.052 | 0.328 | 0.072 | 11.237 | 3 | 0.011 | 0.559 | 73.74 |
|  |  | Ökten et al. | 17 | 0.990 | 0.020 | 34 | 1.000 | 0.010 | -0.700 | 0.305 |  |  |  |  |  |  |  |  |  |  |
|  |  | Brown et al. | 13 | 0.957 | 0.038 | 43 | 0.981 | 0.032 | -0.708 | 0.324 |  |  |  |  |  |  |  |  |  |  |
|  |  | Constantinescu et al. | 40 | 0.960 | 0.046 | 17 | 0.950 | 0.023 | 0.243 | 0.290 |  |  |  |  |  |  |  |  |  |  |
| L2D:4D | Male | Ökten et al. | 9 | 0.980 | 0.030 | 18 | 1.00 | 0.030 | -0.646 | 0.418 | -0.218 | -0.660 | 0.224 | 0.226 | 0.334 | 5.648 | 3 | 0.130 | 0.303 | 45.66 |
|  |  | Brown et al. | 16 | 0.931 | 0.035 | 28 | 0.955 | 0.039 | -0.627 | 0.321 |  |  |  |  |  |  |  |  |  |  |
|  |  | Constantinescu et al. | 24 | 0.959 | 0.035 | 7 | 0.943 | 0.055 | 0.390 | 0.433 |  |  |  |  |  |  |  |  |  |  |
|  |  | Nave et al. | 45 | 0.913 | 0.023 | 39 | 0.913 | 0.020 | 0.000 | 0.219 |  |  |  |  |  |  |  |  |  |  |
|  | Female | Ciumas et al | 11 | 0.979 | 0.027 | 13 | 1.005 | 0.033 | -0.825 | 0.428 | -0.245 | -0.452 | -0.039 | 0.105 | 0.020 | 4.406 | 5 | 0.493 | 0 | 0 |
|  |  | Buck et al. | 66 | 0.925 | 0.021 | 69 | 0.927 | 0.029 | -0.078 | 0.172 |  |  |  |  |  |  |  |  |  |  |
|  |  | Ökten et al. | 17 | 0.990 | 0.040 | 34 | 0.990 | 0.020 | 0 | 0.297 |  |  |  |  |  |  |  |  |  |  |
|  |  | Brown et al. | 13 | 0.952 | 0.025 | 43 | 0.968 | 0.032 | -0.516 | 0.320 |  |  |  |  |  |  |  |  |  |  |
|  |  | Constantinescu et al. | 40 | 0.944 | 0.036 | 17 | 0.955 | 0.028 | -0.320 | 0.291 |  |  |  |  |  |  |  |  |  |  |
|  |  | Nave et al. | 45 | 0.917 | 0.023 | 31 | 0.925 | 0.024 | -0.338 | 0.235 |  |  |  |  |  |  |  |  |  |  |
| M2D:4D | Male | Brown et al. | 16 | 0.934 | 0.037 | 28 | 0.957 | 0.035 | -0.632 | 0.321 | -0.474 | -0.978 | 0.030 | 0.257 | 0.065 | 0.613 | 1 | 0.434 | 0 | 0 |
|  |  | Constantinescu et al. | 24 | 0.950 | 0.028 | 7 | 0.956 | 0.031 | -0.204 | 0.430 |  |  |  |  |  |  |  |  |  |  |
|  | Female | Brown et al. | 13 | 0.954 | 0.026 | 44 | 0.975 | 0.030 | -0.710 | 0.323 | -0.329 | -1.001 | 0.343 | 0.343 | 0.338 | 2.511 | 1 | 0.113 | 0.376 | 60.17 |
|  |  | Constantinescu et al. | 40 | 0.952 | 0.035 | 17 | 0.952 | 0.022 | 0 | 0.290 |  |  |  |  |  |  |  |  |  |  |
| D<sub>[R-L]</sub> | Male | Brown et al. | 16 | 0.006 | 0.033 | 28 | 0.003 | 0.028 | 0.099 | 0.314 | -0.278 | -1.112 | 0.556 | 0.425 | 0.513 | 2.520 | 1 | 0.112 | 0.471 | 60.32 |
|  |  | Constantinescu et al. | 24 | -0.018 | 0.053 | 7 | 0.027 | 0.072 | -0.763 | 0.441 |  |  |  |  |  |  |  |  |  |  |
|  | Female | Brown et al. | 13 | 0.005 | 0.037 | 44 | 0.013 | 0.024 | -0.288 | 0.317 | 0.146 | -0.674 | 0.966 | 0.418 | 0.728 | 3.748 | 1 | 0.053 | 0.507 | 73.32 |
|  |  | Constantinescu et al. | 40 | 0.016 | 0.041 | 17 | -0.005 | 0.026 | 0.556 | 0.294 |  |  |  |  |  |  |  |  |  |  |

**Table 6.** Comparison of meta-analytic effect sizes observed by Hönekopp and Watson (2010) and by the current study

|  | Hönekopp & Watson (2010) |  | Current study |  | Percentage change |
| --- | --- | --- | --- | --- | --- |
|  | *p* | *d* | *p* | *g* |  |
| **Male R2D:4D** | 0.061 | 0.94 | 0.019 | 0.513 | -54.57% |
| **Male L2D:4D** | 0.013 | 0.63 | 0.334 | 0.218 | -34.60% |
| **Female R2D:4D** | < 0.001 | 0.91 | 0.072 | 0.591 | -64.95% |
| **Female L2D:4D** | 0.007 | 0.75 | 0.02 | 0.245 | -32.67% |
| **Mean effect size** |  | 0.81 |  | 0.392 | -46.70% |

*Note.* We compare here the original effect size estimates reported by Hönekopp and Watson (2010) (*d*) with those observed for the current study (*g*); both *d* and *g* indicate standardised mean difference, and the difference between these metrics is negligible.

**Figure 3.**
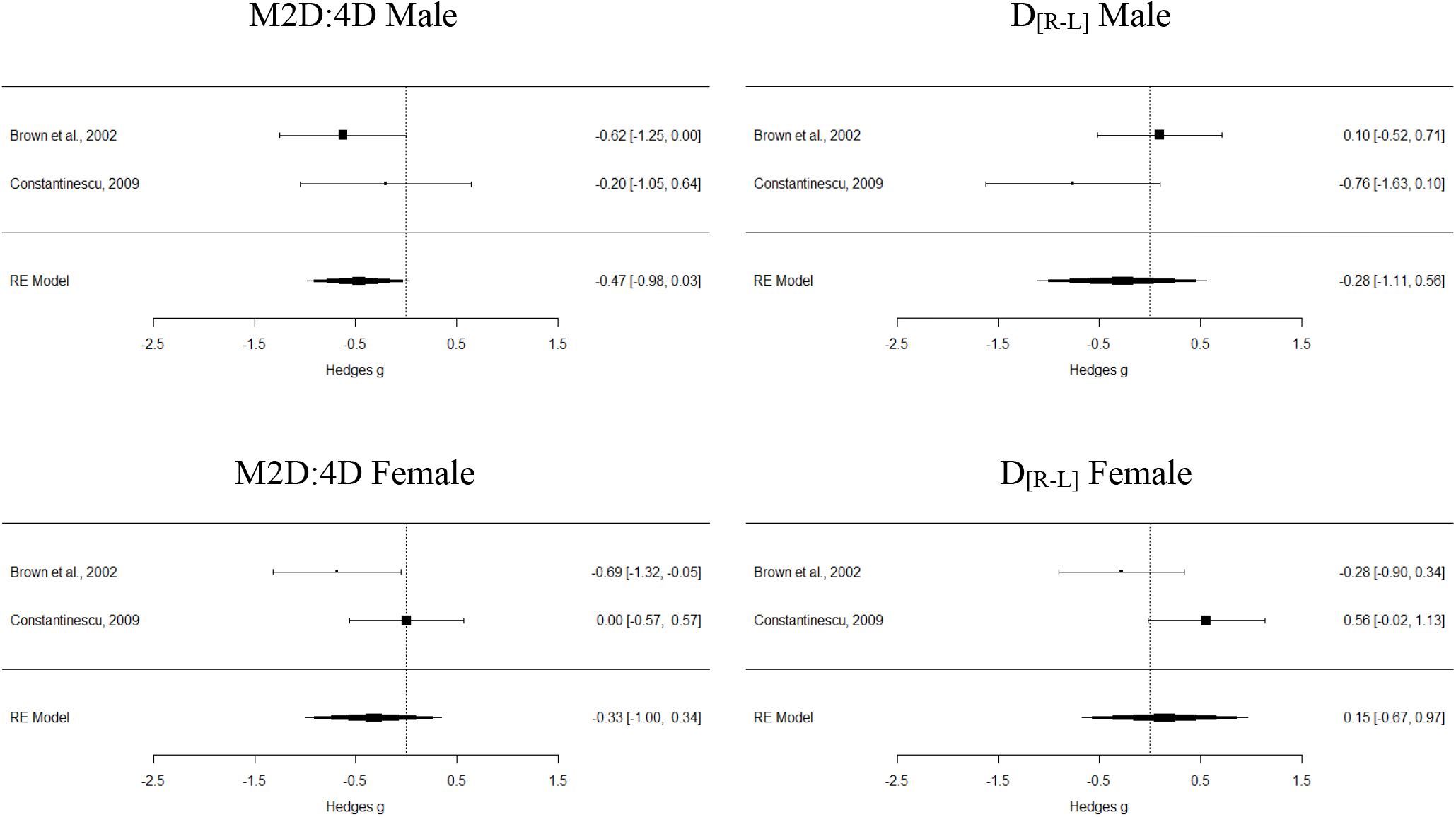
Forest plot summary for each meta-analysis comparing aggregated 2D:4D measures for individuals with CAH to controls for each sex.

**Figure 4.**
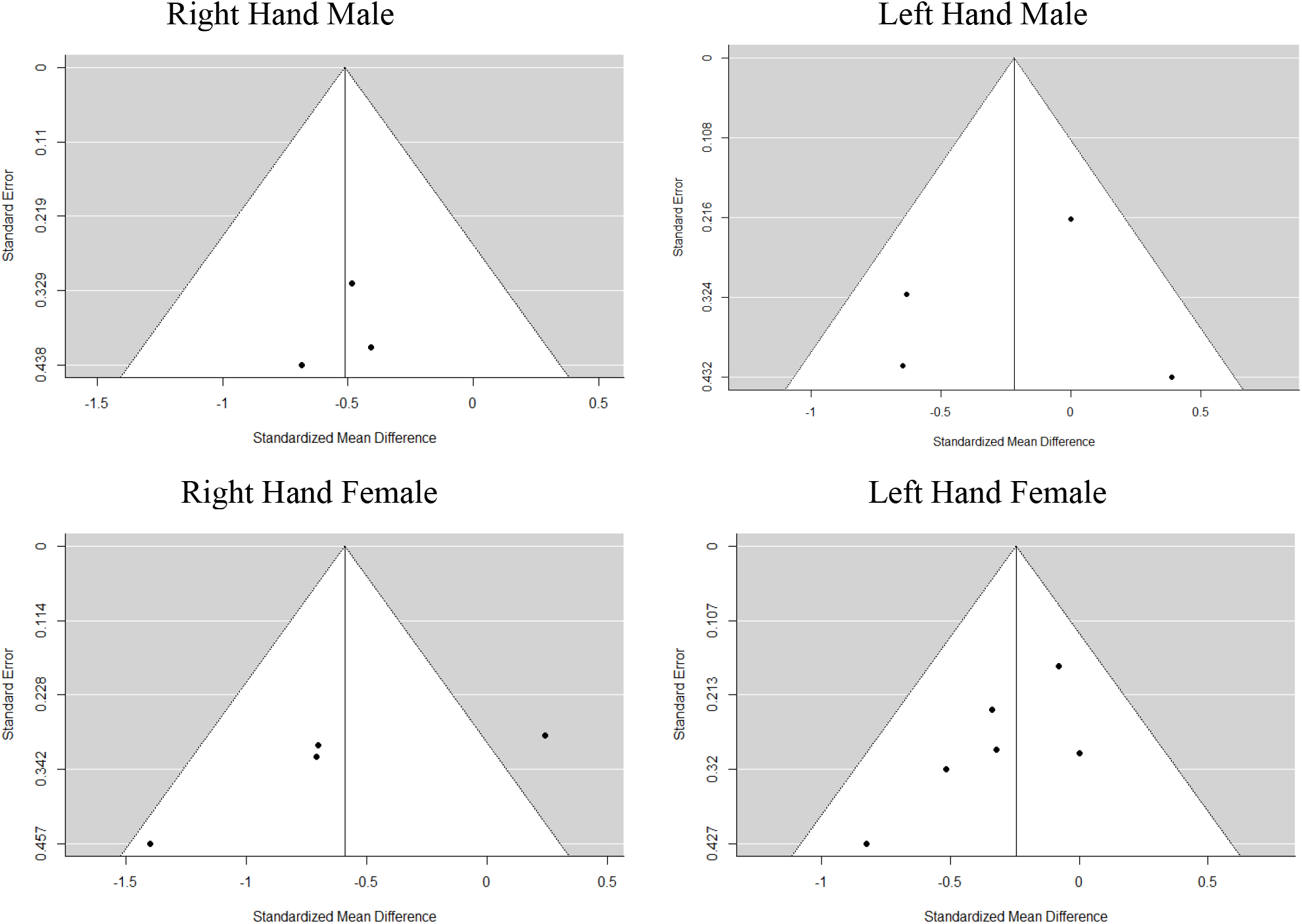
Funnel plots for each meta-analysis comparing mean 2D:4D for individuals with CAH to controls for each sex and hand combination.

We also conducted a meta-analysis that included only studies that used radiographs to measure 2D:4D. It was only possible to perform meta-analyses on the left hand, as only Ökten et al. (2002) used radiographs on the right hand. Ökten et al. (2002) and Nave et al. (2020) used radiographs on the left hand of males. The difference between male participants with and without CAH remained non-significant for the left hand (*g*[95%CI] = −0.225 [-0.829, 0.378], *SE* = 0.308, *p* = 0.465, *Q*(1) = 1.880, *p* = 0.170, τ = 0.313, *I*^2^ = 46.81). For females, Ökten et al. (2002), Buck et al. (2003) and Nave et al. (2020) used radiographs on the left hand. Here the difference was no longer statistically significant between control and CAH participants (*g*[95%CI] = −0.139 [-0.385, 0.108], *SE* = 0.126, *p* = 0.270, *Q*(2) = 1.062, *p* = 0.588, τ = 0, *I*^2^ = 0).

The data provided by Nave et al. (2020) also presented a complication, in that they incorporated multiple cases where participants had 2D:4D measured on multiple occasions. We therefore performed three meta-analyses. The data we presented above take the mean measure for each individual participant across all measures taken for that individual. However, we also performed meta-analyses which took the value from only the first and then only the last measure from each participant. Taking the first value provided results that contrast with the analysis using mean measures as both male and female comparisons identified significant differences in 2D:4D between CAH and non-CAH samples (L2D:4D males: *g*[95%CI] = −0.363 [−0.668, −0.059], *SE* = 0.155, *p* = 0.020, *Q*(3) = 4.199, *p* = 0.241, τ < 0.001, *I*^2^ = 0; 2D:4D females: *g*[95%CI] = −0.302 [−0.535, −0.068], *SE* = 0.119, *p* = 0.011, *Q*(5) = 5.62, *p* = 0.345, τ = 0.120, *I*^2^ = 16.55). However, taking the final measure provided results that align with using the average measure in that male L2D:4D difference was not statistically significant (*g*[95%CI] = − 0.162 [−0.682, −0.348], *SE* = 0.260, *p* = 0.533, *Q*(3) = 7.361, *p* = 0.061, τ = 0.394, *I*^2^ = 58.67), while the female L2D:4D difference remained statistically significant (*g*[95%CI] = −0.228 [−0.434, −0.022], *SE* = 0.105, *p* = 0.031, *Q*(5) = 4.225, *p* = 0.518, τ = 0.001, *I*^2^ = 0). These findings may point towards the importance of considering differences in bone maturation between males and females with CAH and male and female controls.

#### Male vs Female CAH Meta-Analysis Results

In addition to our pre-registered analysis comparing those with and without CAH within sex, we also present exploratory analyses which compare 2D:4D between male and female participants with CAH. As in section 4.1, we summarise the values in Forest Plots (**Figure 5**) and provide greater detail in **Table 7**. Only the difference between R2D:4D for male and female patients with CAH was statistically significant. Egger’s test of small study effects did not identify statistically significant effects (R2D:4D: *z* = −0.023, *p* = 0.982; L2D:4D: z = −0.850, *p* = 0.396). However, the low number of studies provided low power for this test and prevented its calculation for M2D:4D and D_[R-L]_.

**Table 7.** Summary of meta analyses of the difference between 2D:4D for male and female participants with CAH.

| Digit Ratio | Study | Male CAH |  |  | Female CAH |  |  | Effect Size |  | *g* | Meta-Analyses (95%CI) |  |  |  | Q | Heterogeneity |  |  |  |
| --- | --- | --- | --- | --- | --- | --- | --- | --- | --- | --- | --- | --- | --- | --- | --- | --- | --- | --- | --- |
| | | *n* | *M* | *SD* | *n* | *M* | *SD* | *g* | *SE* | | LCI | UCI | *SE* | *p* | | (df) | *p* | $\tau$ | *I*<sup>2</sup> |
| **R2D:4D** | Ökten et al. | 9 | 0.98 | 0.03 | 17 | 0.99 | 0.02 | -0.407 | 0.416 | -0.429 | -0.803 | -0.055 | 0.191 | 0.025 | 0.012 | 2 | 0.994 | 0 | 0 |
|  | Brown et al. | 16 | 0.937 | 0.045 | 13 | 0.957 | 0.038 | -0.463 | 0.379 |  |  |  |  |  |  |  |  |  |  |
|  | Constantinescu et al. | 24 | 0.941 | 0.042 | 40 | 0.960 | 0.046 | -0.421 | 0.261 |  |  |  |  |  |  |  |  |  |  |
| **L2D:4D** | Ökten et al. | 9 | 0.98 | 0.03 | 17 | 0.99 | 0.04 | -0.262 | 0.414 | -0.118 | -0.557 | 0.320 | 0.224 | 0.598 | 6.387 | 3 | 0.094 | 0.325 | 54.26 |
|  | Brown et al. | 16 | 0.931 | 0.035 | 13 | 0.952 | 0.025 | -0.659 | 0.384 |  |  |  |  |  |  |  |  |  |  |
|  | Constantinescu et al. | 24 | 0.959 | 0.035 | 40 | 0.944 | 0.036 | 0.416 | 0.261 |  |  |  |  |  |  |  |  |  |  |
|  | Nave et al. | 45 | 0.913 | 0.023 | 45 | 0.917 | 0.023 | -0.172 | 0.211 |  |  |  |  |  |  |  |  |  |  |
| **M2D:4D** | Brown et al. | 16 | 0.934 | 0.037 | 13 | 0.954 | 0.026 | -0.597 | 0.382 | -0.255 | -0.760 | 0.250 | 0.258 | 0.322 | 1.354 | 1 | 0.245 | 0.194 | 26.15 |
|  | Constantinescu et al. | 24 | 0.950 | 0.028 | 40 | 0.952 | 0.035 | -0.061 | 0.258 |  |  |  |  |  |  |  |  |  |  |
| **D<sub>[R-L]</sub>** | Brown et al. | 16 | 0.006 | 0.033 | 13 | 0.005 | 0.037 | 0.028 | 0.373 | -0.393 | -1.142 | 0.356 | 0.382 | 0.304 | 2.814 | 1 | 0.093 | 0.437 | 64.46 |
|  | Constantinescu et al. | 24 | -0.018 | 0.053 | 40 | 0.016 | 0.041 | -0.733 | 0.266 |  |  |  |  |  |  |  |  |  |  |

**Figure 5.**
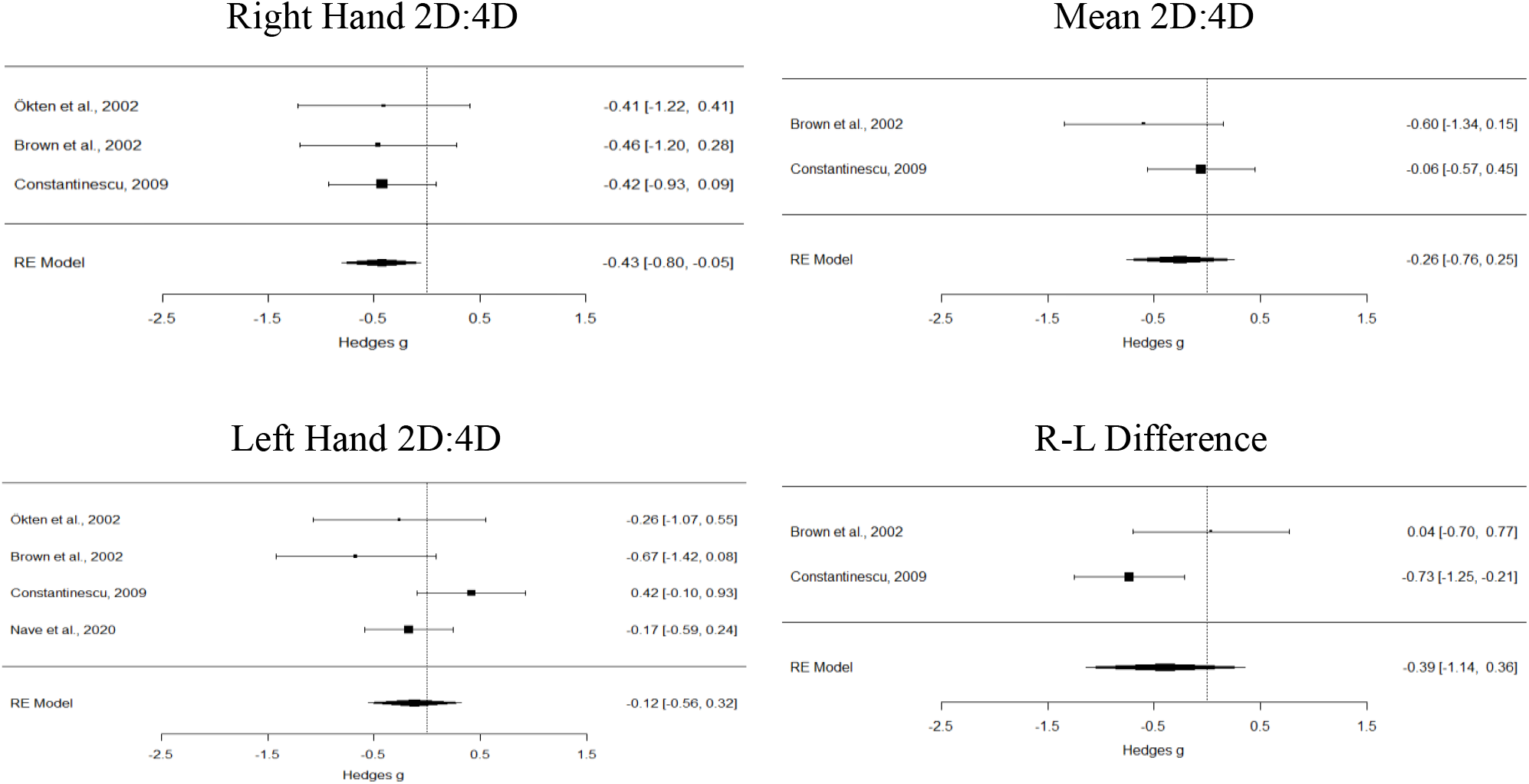
Forest plot summary for each meta-analysis comparing 2D:4D measures between males and females with CAH.

## Discussion

The present study used a systematic literature search to identify as close as possible all studies that have examined 2D:4D in people with CAH. We identified 12 articles relating to nine studies, eight of which reported comparisons between CAH cases and controls. The main findings from the systematic review are that: (1) relatively little research in this area has been published since the meta-analysis of Hönekopp and Watson (2010), (2) most studies have examined small samples, (3) research has been heterogeneous in terms of sample size, country of origin, age-range of participants, type of control group employed, and method used for measuring digit ratio, (4) no studies have specifically examined 2D:4D in CAH caused by enzyme deficiencies other than 21-hydroxylase, (5) no studies have specifically examined 2D:4D in non-classical (i.e. late-onset) CAH samples, (6) only one study has examined differences in 2D:4D between patients with salt-wasting and simple virilising forms of classical CAH, and (7) 2D:4D in CAH samples may increase during childhood similar to controls.

Meta-analyses showed that digit ratios were lower in CAH cases compared with controls for each sex/hand combination, although the effects for R2D:4D in females (*p* = 0.072) and L2D:4D in males (*p* = 0.334) were not statistically significant. Furthermore, if Bonferroni adjustment were employed, the remaining effects (R2D:4D in males, *p* = 0.019; L2D:4D in females, *p* = 0.020) would only retain the required α level of *p* < 0.013 to be considered statistically significant if one-tailed tests were used. Interestingly, our pattern of results was slightly different from that of the meta-analysis by Hönekopp and Watson (2010), in which significant effects were observed for each sex and hand combination other than R2D:4D in males (*p* = 0.061). The addition of new studies has also noticeably reduced the average effect size observed between the previous meta-analysis and the current study, a finding that interestingly appears to mirror that of two meta-analyses of CAH and spatial skills conducted over a very similar same time-period (Collaer & Hines, 2020; cf. Puts et al., 2008). In Hönekopp and Watson (2010) the standardised mean difference (Cohen’s *d*) ranged between 0.63 and 0.94, whereas we report standardised mean differences (Hedge’s *g*) between 0.218 and 0.591 (average reduction in effect size = 46.70%). Part of this reduction could be explained by our use of Hedge’s *g* over Cohen’s *d*, which produces less biased estimates when studies have small samples. However the difference between *d* and *g* is negligible, so it does appear that newer studies have produced smaller estimates of difference. Another potential explanation for the disparity in findings between our study and that of Hönekopp and Watson (2010) is that the latter appears to have treated the infant and young toddler sample of Ökten et al. (2002) as independent from their larger sample. As these samples appear unlikely to have been independent (i.e. although not entirely clear within the article, the smaller sample appears to be comprised of participants from the larger sample), they should not have been included in the same meta-analysis. Doing so is problematic, as it means the same data will be counted twice, likely artificially lowering the standard error of the estimate, which could potentially account for the significant *p* values.

Although we did find evidence to suggest M2D:4D is lower in males and females with CAH relative to male and female controls when re-examining the original data of Brown et al. (2002), these effects were not replicated when reanalysing data from the larger cohort studied by Constantinescu (2009). Further, the meta-analysis combining these estimates found only a marginally significant effect for males (M2D:4D, *p* = 0.065) and no effect for females (*p* = 0.338). No reliable differences between CAH cases and controls were observed for D_[R-L]_ (males: *p* = 0.513; females: *p* = 0.728), casting further doubt on the utility of this variable as an indicator of prenatal sex hormone exposure (Richards et al., 2018).

It appears that digit ratios are typically lower (i.e. more ‘male typical’) in CAH populations than in sex-matched controls ^2^. However, although this might be explainable in terms of the elevated prenatal androgen exposure that characterises CAH, there could also be other explanations. For instance, CAH is additionally associated with reduced concentrations of glucocorticoids and mineralocorticoids, both of which play important roles in bone growth. It was therefore interesting to note that all three studies (Buck et al., 2003; Nave et al., 2020; Ökten et al., 2002) that measured 2D:4D from X-rays reported no significant differences between cases and controls. This could suggest that any difference in 2D:4D between patients with CAH and controls relies on soft tissue rather than bone length, which is consistent with Wallen’s (2009) suggestion that the sex difference in 2D:4D may be due to sex differences in the deposition of adipose tissue in the fingers. However, although Ökten et al. (2002) claimed to have observed significant effects only when examining 2D:4D measured from photocopies (i.e. not when examining X-rays of the same participants), doubt is cast on this premise. This is because our re-analysis of the data reported in Table 3 of that paper (p. 51) revealed that phalangeal R2D:4D was actually significantly lower in females with CAH than in female controls (*p* = 0.021). Further, meta-analyses of the subset of studies that measured L2D:4D from radiographs showed either significant differences in the expected direction or non-significant differences depending on how the data from Nave et al. (2020) were coded. The ambiguity of these findings suggests that further studies comparing radiographic 2D:4D between patients with CAH and controls will be required for firm conclusions to be drawn.

Another important consideration is that classical CAH is characterised by very low gestational cortisol levels, and that this is typically treated by administration of glucocorticoids and mineralocorticoids starting shortly after birth. As sexual differentiation of digit ratios appears prenatally (Galis, Ten Broek, Van Dongen, & Wijnaendts, 2010; Malas, Dogan, Evcil, & Desdicioglu, 2006) yet 2D:4D remains somewhat labile during early infancy (Knickmeyer, Woolson, Hamer, Konneker, & Gilmore, 2011), it is feasible that either prenatal cortisol deficiency and/or early postnatal hormone replacement could affect its development. Although no published studies have examined prenatal or early postnatal cortisol concentrations in relation to 2D:4D in humans, foetal testosterone and cortisol have been shown to be positively correlated (Gitau, Adams, Fisk, & Glover, 2005), and an animal study (Lilley, Laaksonen, Huitu, & Helle, 2010) reported an association between maternal corticosterone levels and offspring 2D:4D ratios in field voles. These observations may suggest that further examination of early cortisol exposure is warranted.

It is noteworthy that, unless considering the lower D_[R-L]_ observed in males with CAH in Constantinescu’s (2009) data, none of the individual studies for which sex differences in CAH samples (i.e. specific differences between males with CAH and females with CAH) could be examined (Brown et al., 2002; Constantinescu, 2009; Kim et al., 2017; Nave et al., 2020; Ökten et al., 2002; Rivas et al., 2014) revealed statistically significant effects in the expected direction (i.e. M<F). However, meta-analysis showed that the sex difference for R2D:4D was significant, and the effect size (*g* = 0.429) is very similar to that reported by Hönekopp and Watson (2010) for typically developing samples (*d* = 0.457). Although the effect for L2D:4D was not statistically significant (*g* = 0.118), this is consistent with the smaller effect size associated with this variable (Hönekopp & Watson, 2010; *d* = 0.376).

When interpreting the current findings, it should be noted individual studies have been diverse, coming from several different countries, and have used photocopies (Brown et al., 2002; Ökten et al., 2002), X-rays (Buck et al., 2003; Nave et al., 2020; Ökten et al., 2002), direct measures (Rivas et al., 2014), and a combination of both direct measures and photocopies (Constantinescu et al., 2010; Constantinescu, 2009) to record 2D:4D. This likely contributed to the relatively high heterogeneity observed for some estimates when these effects were subjected to meta-analysis, which raises some doubt as to the precision of the affected estimates. However, the observation of similar sized sex differences for R2D:4D and L2D:4D in CAH samples as in typically developing populations may cast doubt on the premise that the ratio is strongly affected by prenatal testosterone exposure. This could be because, although testosterone levels are elevated in females with CAH, the prenatal levels for males with CAH may not differ markedly from those of typically developing males. One might therefore predict an absent or partially attenuated sex difference within CAH samples.

A particularly interesting finding from the current study was that the right-left difference in 2D:4D (D_[R-L]_), low values of which have been hypothesised to indicate high exposure to foetal testosterone (Manning, 2002; Manning et al., 2014), did not differ between male or female cases and controls. Although a recent study (Baxter, Wood, Witczak, Bales, & Higley, 2019) reported that high levels of maternal urinary testosterone and testosterone-estrone ratio measured during the first trimester of pregnancy predicted low D_[R-L]_ in the offspring of Titi monkeys, these effects did not retain statistical significance once sex had been controlled for as a covariate (see the analyses presented in the online supplementary materials for that paper). Furthermore, the evidence of such a relationship in humans is even less clear. First, if D_[R-L]_ truly does index individual differences in prenatal androgen exposure in humans, it should arguably exhibit marked sexual differentiation. However, from soon after its inception as a proxy for foetal sex hormone levels. Manning (2002, p. 22) reported that “There may indeed be a tendency for low D*r – l* in males and high D*r – l* in females, but the dimorphism is an elusive one.” Findings from the BBC Internet Study (the largest ever study of digit ratio: male R2D:4D n=126,343; female R2D:4D n=113,725; male L2D:4D n=126,092; female L2D:4D n=113,389) later showed that R2D:4D is only negligibly lower than L2D:4D in males (*d* = 0.01) and negligibly higher than L2D:4D in females (*d* = 0.04) (Manning et al., 2007). Even after considering that the reliability of the self-measured digit ratios used in this study is estimated to be 46% that of expert measurements (Hönekopp & Watson, 2010), and that random measurement errors multiply when ratios are calculed (Martin Voracek, Manning, & Dressler, 2007), the size of any true effect would appear to be very small. When viewed in light of a recent study showing that neither testosterone measured from amniotic fluid nor maternal circulation during the second trimester are correlated with D_[R-L]_ in newborn males or females (Richards et al., 2018), doubt is cast on the validity of this measure as an indicator of prenatal androgen action in humans.

A possible limitation of the current research is the ‘file draw problem’ (Lane, Luminet, Nave, & Mikolajczak, 2016; Rosenthal, 1979), by which studies with small sample sizes and significant results may be more likely to be published than small studies will null findings. This is an issue that has already been posited in specific regard to CAH research (Collaer & Hines, 2020). For instance, Hampson (2016, p. 427) noted that their data were ‘an unfortunate example of this phenomenon’, as they were ‘many years old but were not submitted for publication until now due to the lack of significant group differences’. Although we made extensive efforts to locate unpublished data relating to 2D:4D in CAH samples, we were only able to identify one unpublished dataset (Constantinescu et al., 2010; Constantinescu, 2009). Of specific relevance to 2D:4D of course, is that more than one predictor variable (e.g. R2D:4D, L2D:4D, M2D:4D, D_[R-L]_) is often used to simultaneously assess the same hypothesis. This makes detection of publication bias more difficult because such bias is likely based on whether *any* statistically significant effect is reported, not for *which* predictor variable the effect is observed. This could partly mitigate the existence of publication bias, because having some non-significant findings would not be a barrier to publication. However, it also means that while it is possible to fail to detect any evidence of publication bias in meta-analysis due to the presence of non-significant findings, there could be an unknown number of unpublished studies where none of the 2D:4D comparisons were statistically significant.

In addition to CAH, 2D:4D has been examined in a range of conditions associated with atypical androgen activity, such as CAIS (Berenbaum et al., 2009; van Hemmen et al., 2017), cryptorchidism and/or hypospadias (Abbo et al., 2015; Hwang et al., 2014; O’Kelly, DeCotiis, Zu’bi, Farhat, & Koyle, 2020), polycystic ovarian syndrome (PCOS) (Cattrall, Vollenhoven, & Weston, 2005; Lujan, Bloski, Chizen, Lehotay, & Pierson, 2010; Pandit, Setiya, Yadav, & Jehan, 2016; Roy, Kundu, Sengupta, & Hazra, 2016), autism (Hönekopp, 2012; Manning et al., 2001; Schieve et al., 2018; Teatero & Netley, 2013), ADHD (Martel, Gobrogge, Breedlove, & Nigg, 2008), and gender dysphoria/gender identity disorder and gender variance (Richards, Wei, & Hendriks, 2020; Voracek, Kaden, Kossmeier, Pietschnig, & Tran, 2018). Interest has also been expressed in examining digit ratio in populations with sex chromosome aberrations (Voracek & Dressler, 2007, 2009). Although Manning et al. (2013) reported high 2D:4D in males with Klinefelter’s syndrome (47XXY) compared to their unaffected relatives, this effect has not yet been replicated, and, as far as we are aware, no research has yet examined 2D:4D in relation to Jacob’s syndrome (47XYY), Turner’s syndrome (54XO), or triple X syndrome (47XXX). Regarding Turner’s syndrome, Necić & Grant (1978, p. 311) noted that ‘A short 4th metacarpal is one of the “text-book” signs of Turner’s syndrome’, which may imply high (feminine) 2D:4D ratios in this patient group. However, it should also be considered that Turner’s syndrome is associated with skeletal aberrations, including fusions of bones in the hands (Preger, Steinbach, Moskowitz, Scully, & Goldberg, 1968), which could make it difficult to interpret findings relating to 2D:4D. Further, it has been suggested that there is some symptom overlap (short stature, varying degrees of virilization, amenorrhoea, menstrual irregularities, infertility) between Turner’s syndrome and CAH, and that an elevated rate of 21-hydroxylase deficiency occurs within Turner’s syndrome populations (Larizza et al., 1994). These observations may represent important confounds that should be taken into account when examining 2D:4D within such patient groups. Other potential avenues would be to examine 2D:4D in 47XYY males and 47XXX females, as well as in relation to enzymatic disorders such as 5α-reductase deficiency, and 17β-hydroxysteroid dehydrogenase 3 deficiency.

## Conclusions

Findings from the current study indicate that 2D:4D is lower in patients with CAH compared with sex-matched controls; however, the meta-analytic effects reported here suggest that these effects are substantially smaller than estimated by earlier studies. Although a previous meta-analysis (Hönekopp & Watson, 2010) reported larger effect sizes, the difference in our findings should be considered in light of the fact that we included data from a relatively large study (Constantinescu, 2009) that used an atypical method for quantifying 2D:4D. Nevertheless, the average effect sizes observed for this literature (i.e. the strength of the association between 2D:4D and CAH, a clinical phenotype categorically known to be characterised by elevated prenatal androgen exposure) are small enough to suggest that even studies of 2D:4D that incorporate large samples (i.e. in the hundreds) may be underpowered. We also found no compelling evidence to suggest that the right-left difference in 2D:4D (D_[R-L]_) is significantly different between CAH populations and controls. This is consistent with observations that D_[R-L]_ does not show consistent or large sex differences (Manning et al., 2007), and that it is not correlated with mid-trimester amniotic or maternal circulating testosterone concentrations (Richards et al., 2018).

## Acknowledgements

The authors would like to thank Prof S. Marc Breedlove for kindly sharing with us the data from the Brown et al. (2002) study, Prof Melissa Hines and Dr Debra Spencer for sharing with us the data from Constantinescu (2009), and each of the authors who responded to our request to identify relevant published and unpublished datasets in this area.

## Footnotes

1 The data for Nave et al. were acquired prior to publication of the current article; the current review article was submitted simultaneously with the manuscript that presents Nave et al.’s empirical study.

2 However, it should be noted that the meta-analysis determined that D_[R-L]_ was actually slightly (but not significantly) higher in females with CAH relative to female controls.

